# Synchronization of Circadian Clock Gene Expression in *Arabidopsis* and *Hyaloperonospora arabidopsidis* and its Impact on Host-Pathogen Interactions

**DOI:** 10.1101/2024.03.27.586998

**Authors:** Osman Telli, Deniz Göl, Weibo Jin, Birsen Cevher-Keskin, Yiguo Hong, John M. McDowell, David J. Studholme, Mahmut Tör

## Abstract

Organisms across all kingdoms have an internal circadian clock running in 24h cycles. This clock affects a variety of processes, including innate immunity in plants. However, the role of pathogen circadian clocks had not been extensively explored. We previously showed that light can influence infection of the oomycete *Hyaloperonospora arabidopsidis* (*Hpa*, downy mildew disease) on its natural host *Arabidopsis thaliana*. Here, we identified *Hpa* orthologs of known circadian clock genes (CCGs) *Drosophila TIMELESS (TIM)* and *Arabidopsis Sensitive to Red Light Reduced 1 (AtSRR1)* genes. Expression of both *HpaTIM* and *HpaSRR1* showed a circadian rhythm when *Hpa* was exposed to constant light. Contrastingly, these two genes were negatively regulated by constant dark exposure. Furthermore, the expression patterns of *HpaTIM* and *HpaSRR1* correlate with those of *AtCCA1* and *AtLHY*, indicating a synchronisation of biological clock genes between the host and the pathogen. In addition, screening mutants of *Arabidopsis* Clock Regulated Genes (*AtCRGs*) with three virulent *Hpa* isolates revealed that mutations in *AtCRGs* influenced *HpaTIM* and *HpaSRR1* expression and *Hpa* development, indicating a functional link between the plant biological clock and virulence. Moreover, sporulation of *Hpa* was reduced by targeting *HpaTIM* and *HpaSRR1* with short synthesized small interfering RNAs, indicating that the pathogen clock is also relevant to virulence. We propose that plant and pathogen clocks are synchronized during infection and that proper regulation of both clocks are genetically necessary for pathogen virulence.

## Introduction

Circadian clocks are endogenous subcellular machines that allow organisms to anticipate predictable environmental changes. Changes in the expression of circadian clock genes (CCGs) contribute to plant adaptation to environmental changes, including growth regulation, photoperiodic control of flowering, and responses to biotic and abiotic stresses (Creux & Harmer, 2019; Luklova *et al*., 2019). Similarly, in fungi such as *Neurospora crassa* and the plant-pathogen *Botrytis cinerea*, circadian rhythms affect many processes, such as nutrient uptake, metabolism, conidial production and virulence (Baker *et al*., 2012; Hevia *et al*., 2015). In addition, the circadian rhythms of plants and microbes can influence the timing and outcome of their interactions, including the establishment of beneficial or harmful relationships and defence responses (Bhardwaj *et al*., 2011; Hubbard *et al*., 2018; Newman *et al*., 2022).

The main external signals that affect circadian regulation are light and temperature (Annunziata *et al*., 2018; Wang *et al*., 2020). The centre of all known circadian clocks contains at least one internal autonomous circadian oscillator, with positive and negative elements that create automatic regulatory feedback loops (Hevia *et al*., 2015). In most cases, these loops are used to create 24h timing circuits (Hennessey & Field, 1992). Components of these loops can directly or indirectly receive environmental input to allow entrainment of the clock to environmental time and transfer temporal information through output pathways to regulate expression of rhythmic clock-regulated genes (CRGs) and rhythmic biological activities (Panda *et al*., 2002).

Studies with the model plant *Arabidopsis* and its obligate downy mildew pathogen *Hyaloperonospora arabidopsidis* (*Hpa*) have illuminated many aspects of plant-pathogen interactions (Holub, 2007; Herlihy *et al*., 2019). For example, several disease resistance genes against *Hpa* have been molecularly cloned from *Arabidopsis*. The *R*-gene *RPP4* encodes a nucleotide-binding leucine-rich repeat protein with Toll/interleukin-1 receptor domains and provides resistance to isolates *Hpa*-Emoy2 and *Hpa*-Emwa1 (Van Der Biezen *et al*., 2002). Recent studies on this gene identified a link between *R*-gene-mediated defence and the circadian clock of the host plant (Wang *et al*., 2011). This raises the question of whether *Hpa*’s clock influences virulence in compatible host plants, and whether the host and pathogen clocks might influence each other. Recently, we reported that all developmental stages of *Hpa* during the infection cycle (germination, mycelial growth, and sporulation) are subject to photoregulation (Telli *et al*., 2020), prompting a search for oscillator and clock-related genes (CRGs) in *Hpa*.

The TIMELESS (TIM) gene regulate circadian clocks across species (Myers *et al*., 1995; Lee *et al*., 1996; Panda *et al*., 2002). In *Drosophila*, TIM facilitates entrainment to light– dark cycles by undergoing degradation induced by light, allowing adaptation to the 24h environmental cycle (Allada & Chung, 2010; Rothenfluh *et al*., 2000). TIM forms a complex with the Period (PER) protein in the evening that represses clock gene transcription. Degradation of this complex is initiated by a phosphorylation cycle at late night (Rosato & Kyriacou, 2002). TIM homologs in other organisms like mice and humans have varied functions, with roles in embryonic development and cell cycle regulation (Unsal-Kaçmaz *et al*., 2005; Gotter *et al*., 2000; Young & Kay, 2001), indicating evolutionary divergence while maintaining coordination with circadian rhythms.

*SRR1* (*SENSITIVITY TO RED LIGHT REDUCED*) is a functional gene for clock-regulated expression during day–night cycle (Staiger *et al*., 2003). It was first identified in *Arabidopsis* and its orthologs have been discovered in various organisms (Johansson & Staiger, 2014). *Arabidopsis srr1* mutants display some disfunctions in hypocotyl elongation, greening, petiole growth and flowering, indicating *SRR1* is multifunctional in *Arabidopsis* (Staiger *et al*., 2003). The mouse *SRR1* homologue plays a role in circadian rhythms and cell proliferation (Adachi *et al*., 2017). Similarly, the yeast SRR1-like protein BER1 (Benomyl REsistant 1) is involved in microtubule stability and cell proliferation (Fiechter *et al*., 2008).

Considering the effects of light on *Hpa* virulence (Telli *et al*., 2020) and the rich information on circadian control in other organisms, we reasoned that it would be useful to examine circadian clock regulation in *Hpa*. Here, we identified *HpaTIM* and *HpaSRR1* in *Hpa*, and investigated their expression patterns under different light regimes. Additionally, we investigated the expression pattern of three well-characterized *Arabidopsis* circadian clock genes, *CIRCADIAN CLOCK-ASSOCIATED 1* (*CCA1*), *TIMING OF CAB EXPRESSION 1* (*TOC1*) and *LATE ELONGATED HYPOCOTYL* (*LHY*) during infection by *Hpa* under different light regimes. We report here that *HpaTIM* and *HpaSRR1* show rhythmic expression and synchrony with the expression of *CCA1* and *LHY*, and we provide genetic evidence from the host and pathogen that supports the biological relevance of this synchronization.

## Results

### *Hpa* encodes clock-related genes

Until now, no investigation on circadian-related genes has been reported for *Hpa*. We addressed this knowledge gap by searching the *Hpa* genome for homologues of important circadian genes from other organisms, using BLAST and domain-searches using protein domains that are characteristic of the relevant proteins in model organisms. Two putative CRGs were identified in the *Hpa* genome: *Timeless* (*Hpa-G810921*, designated *HpaTIM*) and *Sensitive to Red Light Reduced 1 (Hpa-G801448*, designated *HpaSRR1*). Further bioinformatic analyses and EnsemblProtists gene annotation revealed that *HpaTIM* exists as a single-copy gene with two introns that encodes a predicted protein of 1175 amino acids with a molecular mass of 131.7 kDa. Domain and motif searches of *Hpa*TIM revealed two TIMELESS domains (N35-A592; W688-T751) (Supplemental Figure 1). Amino acid sequences of TIMELESS proteins from various species were aligned (Figure 1A) and a phylogenetic tree was constructed (Figure 1C). *Hpa*TIM showed a high amino-acid identity to TIM proteins from other species (Figures 1A and C). *HpaTIM* orthologues were also found in other oomycete pathogens (Supplemental Figure 2). Published transcriptome data in *Arabidopsis* Col-0 inoculated with the avirulent or virulent *Hpa* isolates Emoy2 or Waco9, respectively (Asai *et al*., 2018) indicates that *HpaTIM* is expressed in spores and during infection (Supplemental Figure 3).

**Figure 1.**
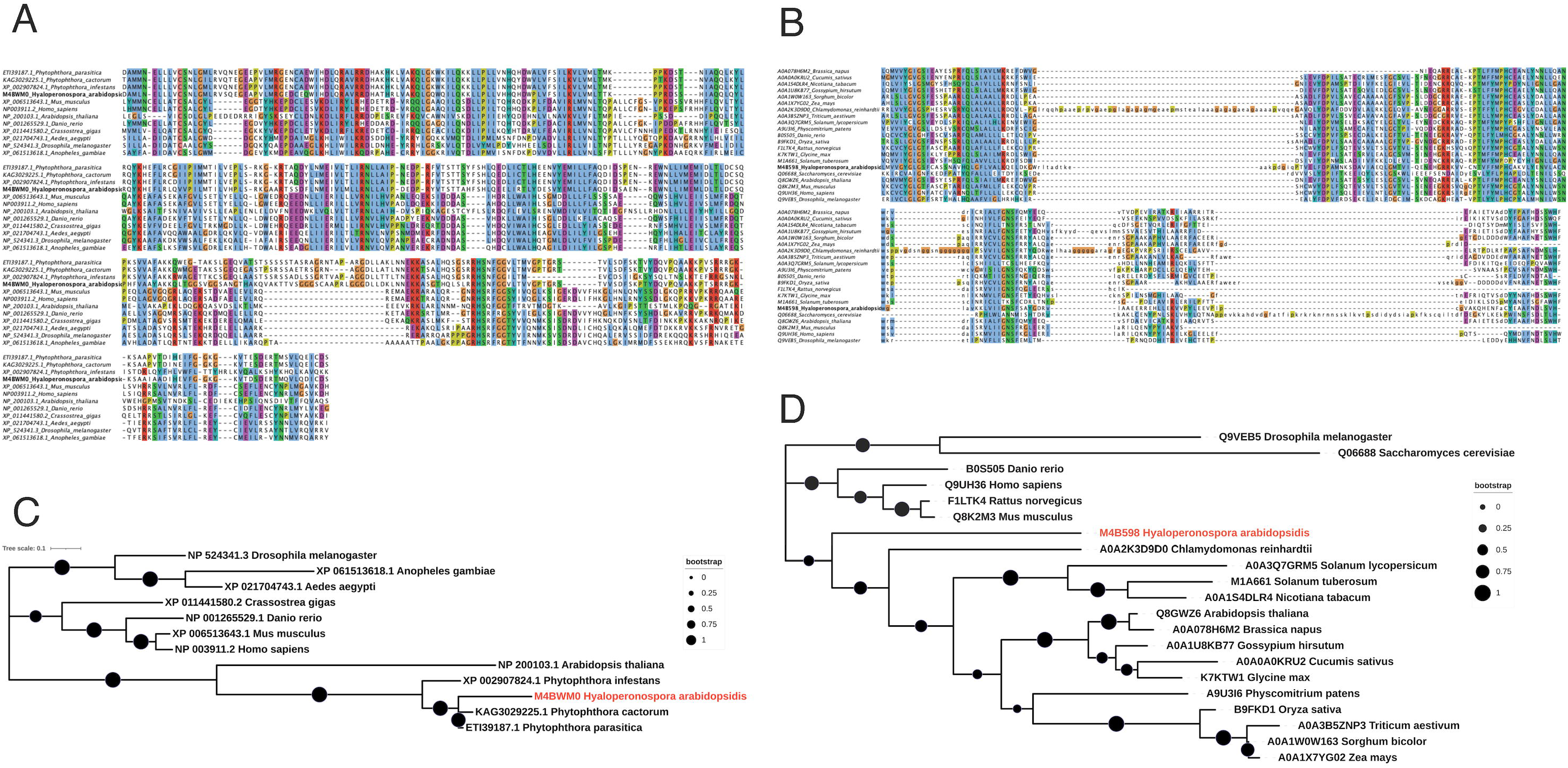
Phylogenies of HpaTIM and Hpa-SSR1. <u>Panel A</u> shows a multiple sequence alignment of the Timeless protein domain (Pfam: PF04821) of HpaTIM against homologues in model organisms. <u>Panel B</u> shows a multiple sequence alignment of the SRR1 domain (Pfam: PF07985) of HpaSRR1 against homologues in plants and model organisms. <u>Panel C</u> shows evolutionary analysis of HpaTIM. <u>Panel D</u> shows the evolutionary history of HpaSRR1. The sizes of the black circles indicate the proportions of 1000 bootstrapped trees in which the associated taxa clustered together. The trees are drawn to scale, with branch lengths measured in the number of substitutions per site.

*HpaSRR1* exists as a single copy in the reference *Hpa* genome and has three predicted introns. The open reading frame of *HpaSRR1* encodes a predicted protein of 335 aa (molecular weight 37.013 kDa). Domain and motif searches revealed a SRR1-like protein domain (H3-S304) (Supplemental Figure 4). Proteins from various species were identified, amino-acid sequences were aligned (Figure 1B) and a phylogenetic tree was constructed (Figure 1D). Additionally, orthologues of HpaSRR1 were identified in other oomycete pathogens, indicating the conserved nature of the gene (Supplemental Figure 5). Expression of this gene was not evident in the published transcriptome data (Asai *et al*., 2018). However, we were able to demonstrate *HpaSSR1* expression during infection, as described in the following section.

### Targeting HpaTIM and HpaSRR1 with small dsRNA reduces Hpa sporulation

Following identification of *HpaTIM* and *Hpa-SSR1*, we used a genetic approach to test whether these genes are necessary for *Hpa* viulence on *Arabidopsis*. *Hpa* is an obligate biotroph and cannot be genetically transformed by conventional approaches. However, a small RNA-based approach was recently developed to reduce the expression of targeted *Hpa* genes through transcriptional or translational silencing (Bilir *et al*., 2019). <u>S</u>hort, <u>s</u>ynthetic, <u>d</u>ouble-<u>s</u>tranded RNAs (SS-dsRNAs) were designed to target *HpaTIM* and *HpaSSR1.* These SS-dsRNAs were mixed with *Hpa*-Emoy2 spores at 5µM concentrations and used to drop-inoculate 7-day old seedings of the disease-susceptible mutant *Arabidopsis* Ws-*eds1*. At 7dpi, plants inoculated with spore suspensions containing 5µM SS-dsRNA targeting *HpaTIM* and *HpaSRR1* showed reduced sporulation (∼50-70%) compared to plants inoculated with untreated spores (Figure 2A). No sporulation was observed with the positive control targeting the essential *Hpa-CesA3* gene as reported before (Bilir *et al.,* 2019). We then checked the relative mRNA abundance of the targeted genes. While there was statistically significant reduction in the expression level of *Hpa-CesA3*, we did not observe any significant decrease in the expression level of *HpaTIM* and *HpaSRR1* (Figure 2B), suggesting that the SS-dsRNAs interfered with translation of *HpaTIM* and *HpaSRR1* rather than transcription.

**Figure 2.**
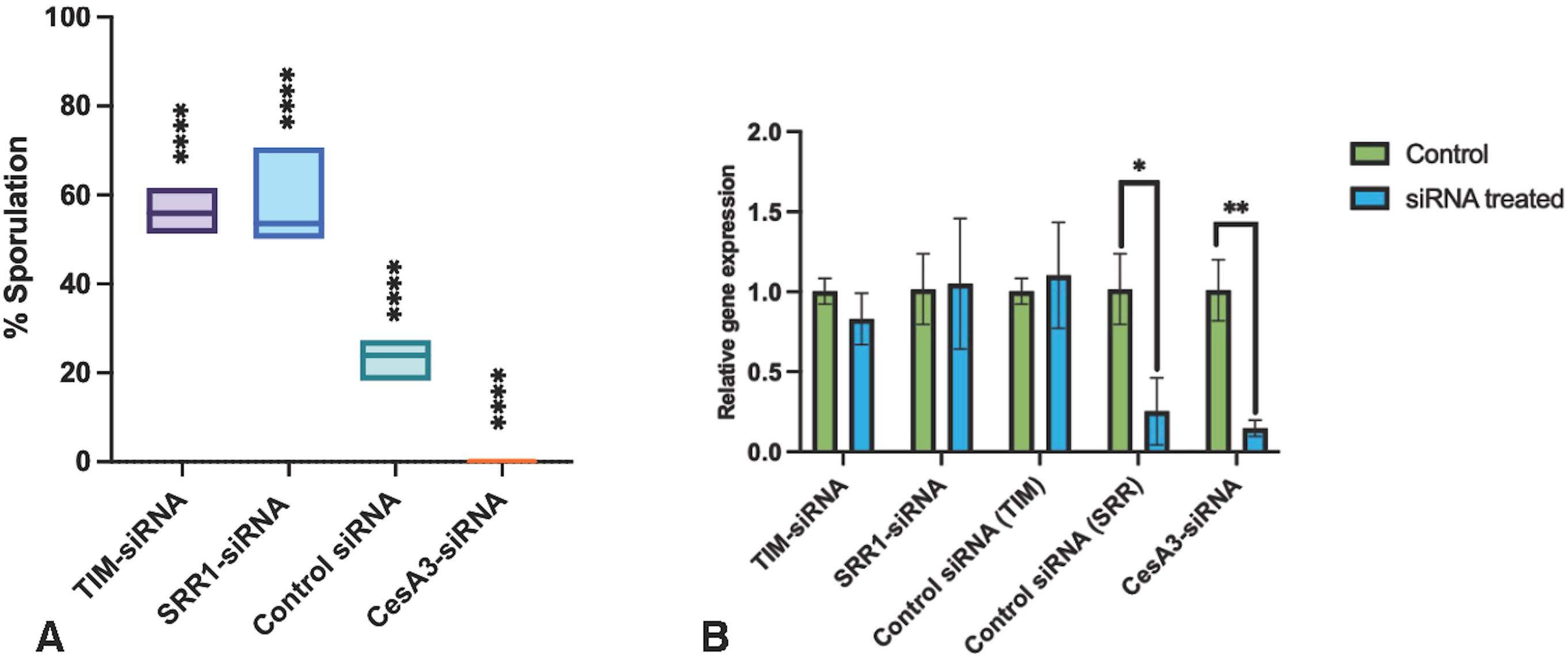
Sporulation of *Hpa* and expression level of *HpaTIM* and *HpaSRR1* targeted with SS-dsRNA. Sporulation was assessed 7dpi **(A)**, and gene expression was determined using qRT-PCR at 3dpi **(B).** *Hpa-CesA3* was targeted as a positive control. An SS-dsRNA that does not inhibit pathogen growth was also used. One-way ANOVA has been performed on data to compare treated samples with control samples. Data expressed as mean ± mean standard deviation. ****p < 0.0001, **p = 0.0039, *p = 0.0102

### *HpaTIM* gene is influenced by light regime

After establishing *HpaTIM* as a strong candidate CRG, with biologically relevant effects on *Hpa* virulence, we wanted to investigate whether *HpaTIM* expression shows a circadian rhythm during infection of Arabidopsis. Because Col-0 is resistant to *Hpa*-Emoy2, we used a susceptible mutant (Col-*rpp4*) seedlings in the experiments. Seven-day old seedlings were infected at 0 ZT hour with *Hpa* spores, allowed to grow under a “normal” light regime (12h D / 12h L) for the first 3 days, and then were exposed to constant light or constant dark for 24h between 3 to 4dpi. After the 24h exposure, samples were taken every 6h from infected plants between 4-7dpi. Similarly, samples were also taken from infected plants between 4-7dpi that were kept under the normal light cycle to serve as controls. Abundance of the *HpaTIM* mRNA was quantified with qRT-PCR.

*HpaTIM* displayed a rhythmic pattern with a 24h period under a normal 12h D / 12h L (DL) cycle (Figure 3A and B). Expression peaked at the beginning of each light cycle (dawn) and gradually decreased until the beginning of the dark cycle (Figure 3A). After 24h constant light exposure (between 3 and 4dpi), the expression pattern of the *HpaTIM* was disrupted slightly: although the amplitude was not changed, the length of the period shortened to around 18h, but was still rhythmic indicating that this gene has a circadian pattern. At 60 hrs after constant light treatment (6 dpi), *HpaTIM* expression returned to its normal 24h-period cycle (Figure 3A).

**Figure 3.**
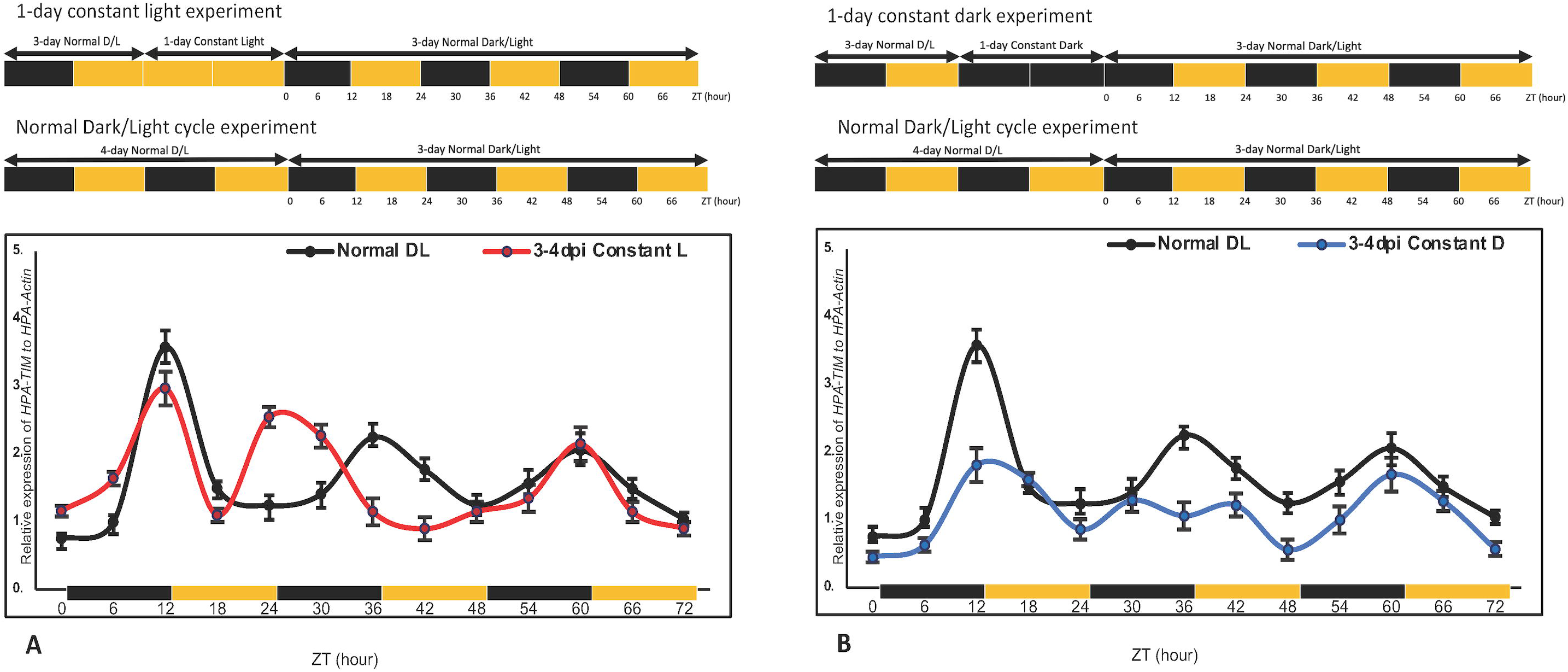
Expression analyses of *HpaTIM* in different light regimes. **A.** Expression of *HpaTIM* gene in a normal light cycle (black line) and constant light exposure between 3dpi to 4dpi followed by a normal light cycle (red line). **B.** Expression pattern of *HpaTIM* normal cycle (black line) and subsequently exposed to constant dark between 3dpi-4dpi followed by normal cycle (blue line). For the first three days post inoculation (dpi), all samples were kept under the 12h Dark/ 12h Light (D/L) cycle, referred to hereafter as “normal”. After 4dpi, ZT time (Zeitgeber Time) started, and samples were taken every 6h over 3 days. Experiments started at dusk. While black blocks represent dark periods, yellow blocks represent light periods. *Hpa-Actin* was used as a standard. The experiment was repeated 3 times and similar results were obtained. Standard error of mean for 3 biological replicas are indicated.

The amplitude of *HpaTIM* expression in tissues exposed to constant dark for 24 hours was much lower compared to tissues exposed to normal or constant light. There was also a shift in the expression cycle of *HpaTIM* in tissues exposed to 24h constant dark between 3 and 4 dpi (Figure 3B). The period was shortened, however, by day 6 the cycle of expression had almost returned to normal.

In a second set of experiments, inoculated plants were allowed to grow for 3 days under a normal light regime (12h D / 12h L) and were then exposed to 4 days constant light or constant dark (Figure 4). The expression of *HpaTIM* under constant light showed reduced period and increased amplitude in comparison to the control. The expression peaked at different times, such as into 4 and 5 dpi, peaks were observed at dusk, however into 6 and 7dpi, peaks were observed at dawn. *HpaTIM* expression under constant dark did not display a proper period and a clear peak (Figure 4), suggesting that it may be totally suppressed.

**Figure 4.**
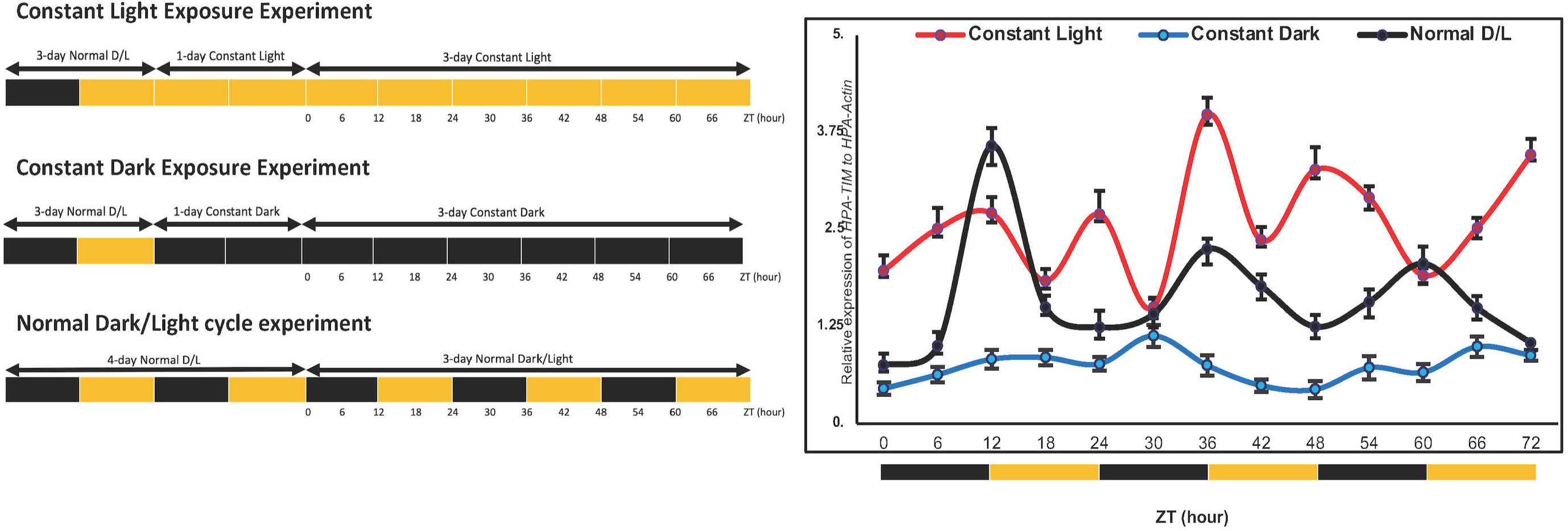
Expression of *HpaTIM* under constant light and constant dark exposure. After an initial 3-day normal D/L conditions, one group was exposed to continuous light for the following 4d while the other group exposed to continuous darkness for the following 4d. Normal D/L cycle time-course served as the control. Samples were taken every 6h and the expression levels of the *HpaTIM* gene were investigated. Experiments started at dusk. While black blocks represent dark periods, yellow blocks represent light periods. *Hpa-Actin* was used as a standard. The experiment was repeated 3 times and similar results were obtained. Standard error of mean for 3 biological replicas are indicated.

### HpaSRR1 shows rhythmic expression

Similar to *HpaTIM*, we investigated *HpaSRR1* expression under different light regimes. Inoculated plants were grown under normal light regimes for 3d (12h D / 12h L) and then were exposed to constant light or constant dark for 24h. Samples were taken every 6h for 3 days and expression pattern of *HpaSRR1* was determined. Under a normal DL regime, expression of *HpaSRR1* showed a periodic cycle similar to that of *HpaTIM*.

Expression peaked at dawn for each day (Figure 5). Exposure to constant light at 3-4 dpi did not change the expression pattern of *HpaSSR1;* the gene exhibited a rhythmic expression pattern under the constant light just as in the normal cycle. The amplitude of *HpaSRR1* expression was slightly higher at the beginning but in general, it was very similar to that of the control (Figure 5A).

**Figure 5.**
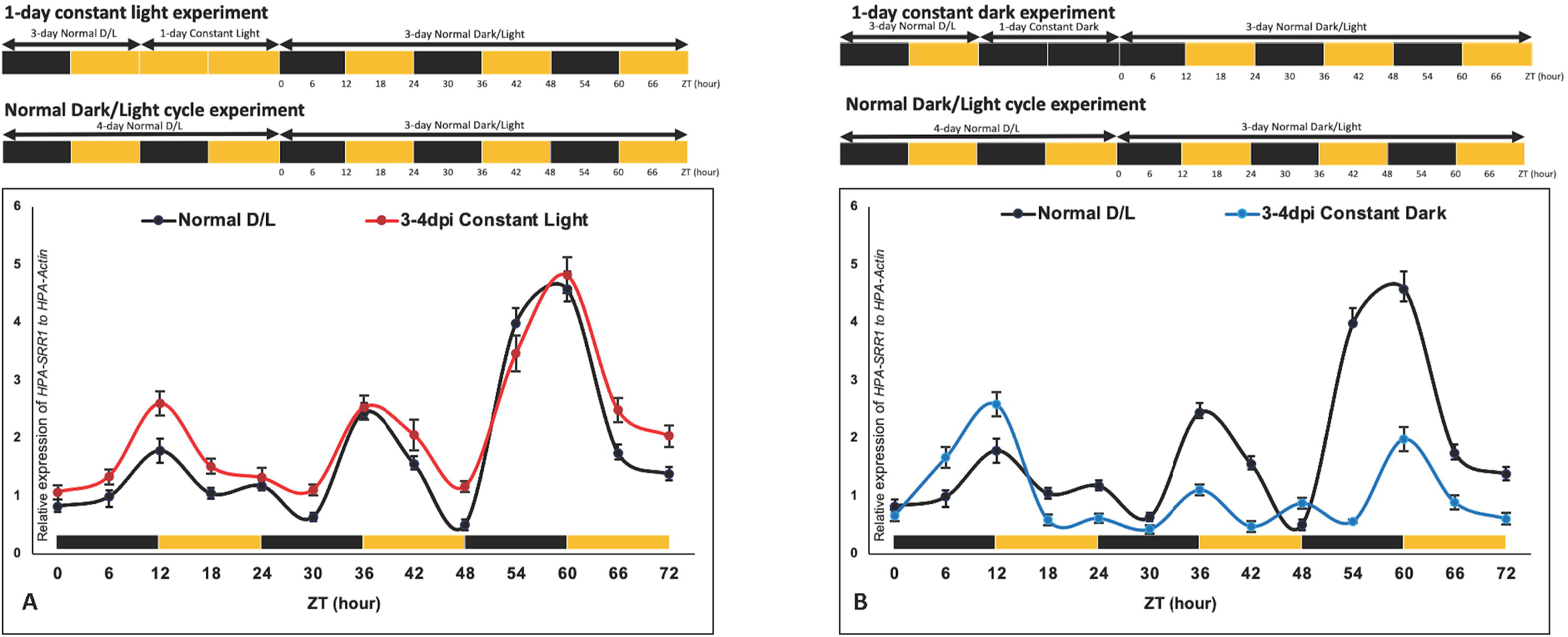
**Expression of *HpaSRR1* in different light regimes**. **A.** Expression of *HpaSRR1* in normal D/L cycle (black line) and then constant light exposure after 3 dpi (blue line). Expression showed rhythmic expression pattern just as observed with a normal cycle. **B.** Expression of *HpaSRR1* in a normal light cycle (black line) followed by exposure to constant dark between 3 dpi to 4 dpi, followed by a normal light cycle (blue line). Experiments started at dusk. Black bars represent dark periods, yellow bars are light periods. *Hpa-Actin* was used as a standard. The experiment was repeated 3 times and similar results were obtained. Standard error of mean for 3 biological replicas are indicated.

*HpaSRR1* expression levels were observed with samples, which were exposed to constant dark at 3-4 dpi (Figure 5B). Similar to the DL and 3-4 dpi constant light series, all peaks in expression levels were observed at dawn for each day. However, expression levels in samples exposed to constant dark were lower than that found with constant light (4d/0h) and 4d/6h. These findings indicate that *HpaSRR1* expression may have rhythmic oscillation, even under the irregular light regime for a short period where expression patterns were not broken or shifted (Figure 5A).

### Arabidopsis and Hpa timing systems are synchronised

*Hpa* has an obligate biotrophic lifestyle and cannot exist apart from its host (Woods-Tör *et al.,* 2018). Given this intimate relationship, it is plausible that the *Hpa*’s clock is synchronized with the *Arabidopsis*’ clock during infection. In the *Arabidopsis* circadian clock system, expression of *CCA1 (Circadian Clock Associated 1), LHY (Late Elongated Hypocotyl)* and *TOC1 (Timing of CAB expression 1*) genes are commonly used as biomarkers for circadian regulation (McClung, 2006). Expression of *CCA1* and *LHY* has been reported to peak at dawn, and *TOC1* expression peaks have been observed at dusk. (McClung, 2006; De Caluwé *et al*., 2016). Reciprocal regulation between *CCA1*, *LHY*, and *TOC1* is thought to provide a feedback loop mechanism which is essential for circadian rhythmicity in *Arabidopsis* (Alabadi *et al*., 2001)

We wanted to determine whether there is a correlation between the expression pattern of *HpaTIM* and *HpaSRR1* genes compared to *Arabidopsis CCA1, LHY* and *TOC1* genes. First, we examined the expression patterns of *CCA1* and *LHY* over 3d between 4 and 7dpi. Expression of both *CCA1* and *LHY* peaked at dawn during this period (Figure 6A). *CCA1* expression levels were higher than that of *LHY.* With *TOC1* expressions, peaks were observed at dusk, and were lowest at dawn (Figure 6A). These results were in agreement with published data from uninfected plants (Alabadi *et al*., 2001). Secondly, we compared the rhythmic expression patterns of *HpaTIM* and *HpaSRR1* with that of *CCA1* and *LHY*. We observed that the expression pattern of *HpaTIM* and *HpaSRR1* were very similar to that of *CCA1* and *LHY* (Figure 6B). In all series, expression levels were peaked at dawn. Expression levels increased during the dark and decreased in light cycle. These findings suggest that *Arabidopsis-Hpa* pathosystem has a synchronised circadian regulation.

**Figure 6.**
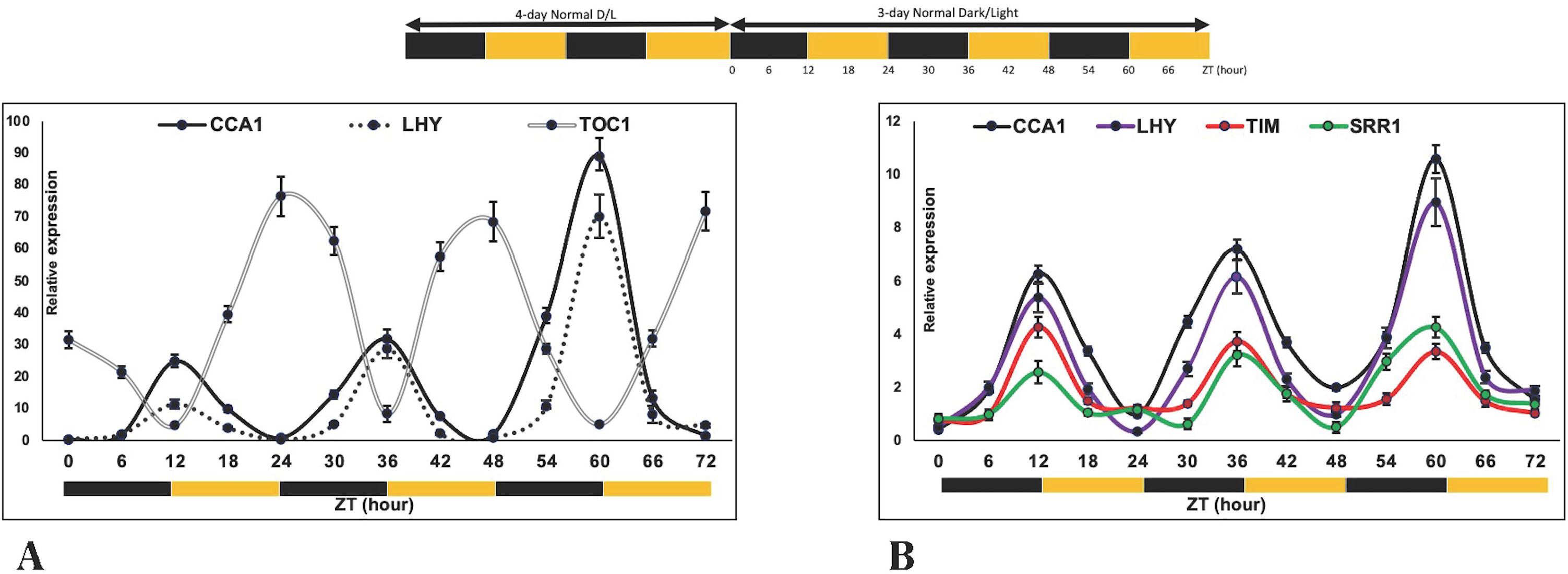
**Expression of *HpaTIM* and *HpaSRR1* compared with *CCA1* and *LHY during infection***. **A)** Expression of *CCA1* (black line), *LHY* (dashed line) and *TOC1* (grey line) in D/L cycle. Samples were taken every 6h into 4 dpi-7 dpi. *At-Actin* was used as a standard. **B)** Comparison of the expression of *HpaTIM* (red line) and *HpaSRR1* (green line) with *CCA1* (black line), *LHY* (purple line). Samples were taken every 6h from 4 dpi-7 dpi. This experiment started at dusk. Black bars represent dark period, yellow bars are light period. Experiment was repeated 3 times and similar results were obtained. Standard error of mean for 3 biological replicas are indicated.

### Mutations in *cca1* and *cca1/lhy* influence *HpaTIM* and *HpaSRR1* expressions

If the *Arabidopsis* and *Hpa* clocks are synchronized, then *Arabidopsis* mutations that affect circadian regulation might disrupt the regulation of CRGs in *Hpa* during the infection cycle. To address this prediction, we investigated expression patterns of *HpaTIM* and *HpaSRR1* during infection of Col-*cca1* single mutants and Col-*cca1*/*lhy* double mutant lines.

*HpaTIM* and *HpaSRR1* expression were assayed between 4 and 6 dpi on the *cca1* mutant line (Figure 7A). Although *HpaTIM* and *HpaSRR1* showed a circadian expression pattern on the Col-*cca1* mutant, this pattern did not exactly match with that of *HpaTIM* and *HpaSRR1* on wild-type Col-0 line (Figure 7A). When compared with the normal pattern; the expression peaks on the *cca1* mutant line were observed not at dawn, but in the middle of the day (4d.18h and 5d.18h), and the lowest points of the expression were observed in the middle of the night (Figure 7A), indicating that the *CCA1* gene influences the timing but not the amplitude of *HpaTIM* and *HpaSRR1* expression.

**Figure 7.**
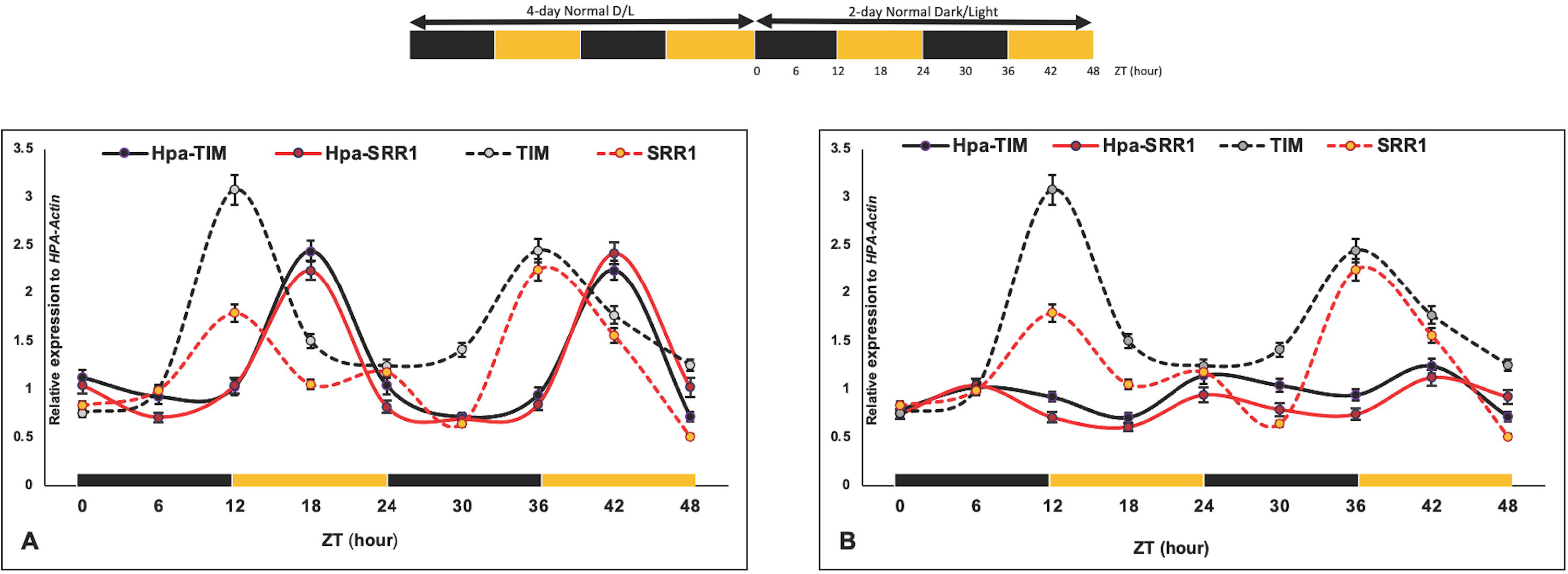
**Expressions of *HpaTIM* and *HpaSRR1* on *Arabidopsis* single or double clock mutants**. Each graph displays expression of these genes in *Hpa* growing on the mutant (solid lines) or wild-type Col-0 (dashed lines). **A)** *HpaTIM* and *HpaSRR1* expressions on Col-*cca1* mutant line. **B)** *HpaTIM* and *HpaSRR1* expression in *cca1/lhy* double mutant line. Col-*cca1* or *cca1/lhy* mutants were inoculated with *Hpa*-Maks9. Samples were taken from inoculated seedlings between 4 and 6 dpi and expression patterns of *HpaTIM* and *HpaSRR1* were quantified with RT-qPCR. Experiments started at dusk. Black bars represent dark periods, yellow bars are light periods. *Hpa-Actin* was used as a normalization control. This experiment repeated 3 times with consistent results. Standard error of mean for 3 biological replicas are indicated.

Similarly, expression levels of *HpaTIM* and *HpaSRR1* were also investigated in the Col-*cca1*/*lhy* double mutants between 4 and 6 dpi for 2d (Figure 7B). The expression levels of *HpaTIM* and *HpaSRR1* observed on the double mutant differed significantly from the expression levels observed on the wild-type Col-0. *HpaTIM* and *HpaSRR1* expression levels were peaked in the middle of the night (4d, 6h), at the beginning of the day (5d) and in the middle of the day (5d,18h) (Figure 7B). The expression patterns of *HpaTIM* and *HpaSRR1* were still similar and parallel to each other, and peaks were shifted (Figure 7B), indicating that *HpaTIM* and *HpaSRR1* are regulated during infection by a common mechanism that requires *Arabidopsis CCA1* and *LHY1* genes.

### Mutation in Arabidopsis CRGs alters *Hpa* sporulation and biomass production

Observation of altered *Hpa* CR gene expression on *Arabidopsis* single and double clock mutants prompted us to determine whether pathogen sporulation and biomass production was affected by mutations in *Arabidopsis* CRGs. We screened 26 single-, 2 double- and 1 triple mutants, along with 3 overexpressors (ox) lines.

Each homozygous mutant line, and *CCA1ox* lines were inoculated with the compatible isolate *Hpa*-Noks1, and the amount of sporulation was calculated and compared to the control Col-0 line (Figure 8A). Overall, 10 lines supported less sporulation than that in the control: *elf3* (75%), *phyb* (75%), *kat2* (73%), *lux* (73%), *cry1* (70%), *lcl5* (65%), *CCA1ox* (60%), *lhy* (60%), *cca1*(47%) and *pif3* (47%) (Figure 8A), indicating that that these CRGs contribute to compatibility. Contrastingly, three mutant lines supported more sporulation compared to the Col-0 control: *prr9* (138%)*, toc1*(135%) and *tic1* (133%) (Figure 8A), suggesting that these genes could be essential for basal defence in *Arabidopsis*.

**Figure 8.**
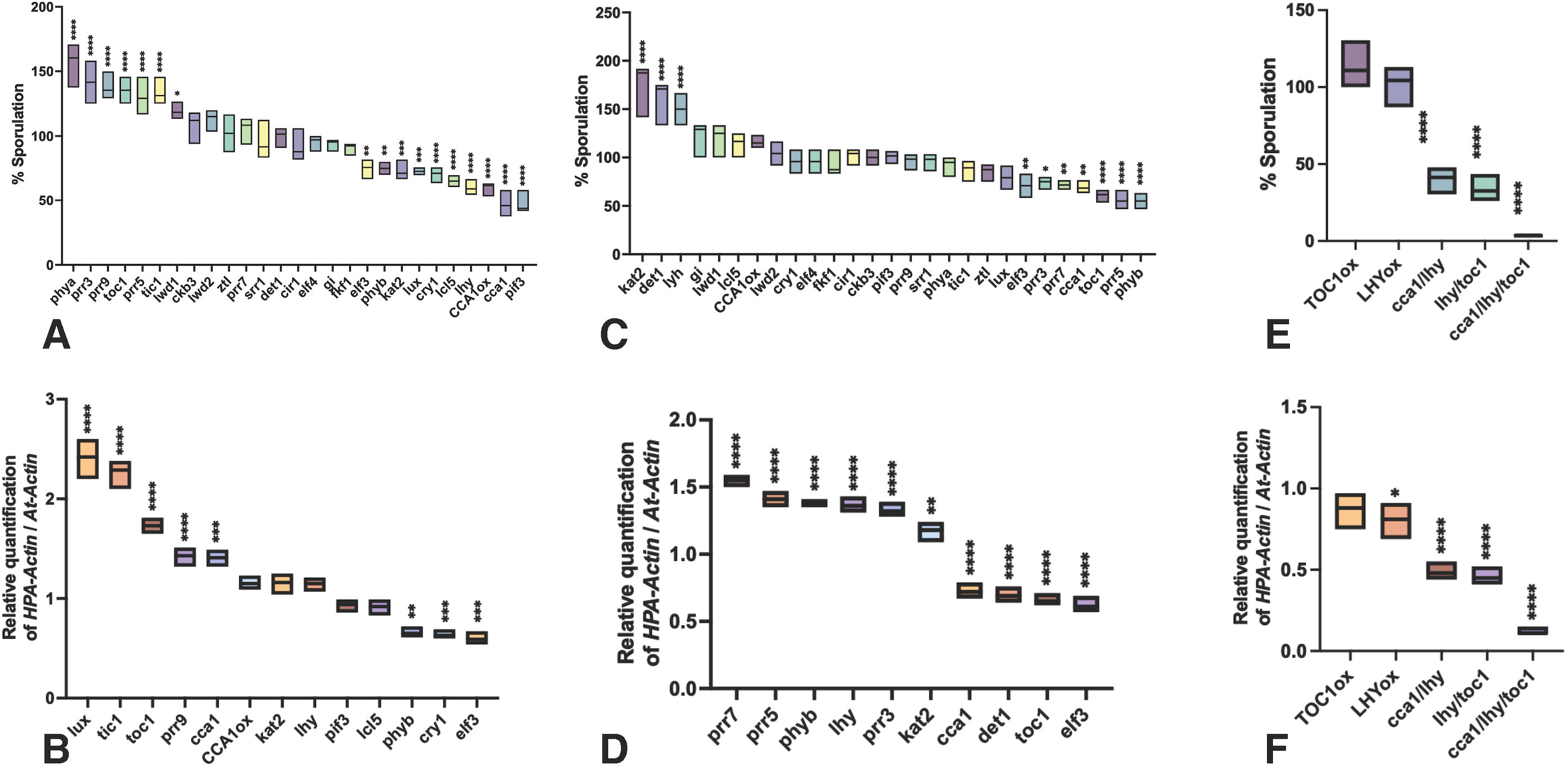
Mutations in *Arabidopsis* clock regulated genes influence sporulation and biomass production. T-DNA null mutants in Col-0 or Ws-0 background and overexpressor (ox) were inoculated with a compatible *Hpa* isolate and the amount of sporulation was calculated at 7 dpi. The biomass of vegetative hyphae was investigated with selected mutant lines 3 dpi using q-PCR. Overexpressors, TOC1ox and LHYox were also included. One-way ANOVA has been performed on data to compare mutant lines with control samples. Data expressed as mean ± mean standard deviation. **A)** Mutant lines inoculated with *Hpa*-Noks1 (CCA1ox, *cca1, pif3, tic1, toc1, prr9, prr5, prr3, phya, lcl5, lhy, cry1* ****p < 0.0001; *lux, kat2*, ***p = 0.0003; *phyb, elf3* **p = 0.0013, *lwd1* *p = 0.0301). **B)** Mutant lines inoculated with *Hpa*-Noks1 (*prr9, lux, toc1, tic1* **** p < 0.0001; *cry1* ***p=0.0007; *elf3* ***p=0.0002; *cca1* ***p=0.0001; *phyb* **p=0.0015). **C)** Mutant lines inoculated with *Hpa*-Maks9 (*toc1, prr5, phyb, lhy, kat2, det1* ****p < 0.0001; *cca1* **p=0.0023; *prr7* **p=0.0069; *elf3* **p=0.0049; *prr3* *p=0.0190). **D)** Mutant lines inoculated with *Hpa*-Maks9 (*det1, elf3, lhy, phyb, prr3, prr7, prr5, toc1, cca1* ****p < 0.0001; *kat2* **p=0.0067). **E)** Mutant lines were inoculated with *Hpa*-Emco5 (*cca1/lhy*, *cca1/lhy/toc1*, *lhy/toc1* ****p < 0.0001). **F)** Mutant lines were inoculated with *Hpa*-Emco5 (*lhy/toc1*, *cca1/lhy/toc1, cca1/lhy* ****p < 0.0001; LHYox *p=0.0232).

The 13 mutant lines displaying a significant sporulation phenotype with *Hpa*-Noks1 were evaluated for vegetative hyphal biomass at 3 dpi. Among the 13 lines, *cry1 phyb* (66%), *cry1* (64%) and *elf3* (60%) produced less biomass production compared to the wild-type control. Contrastingly more biomass was produced in *lux* (241%), *tic1* (226%), toc1(173%), *cca1* (141%) and *prr9* (140%) (Figure 8B). Interestingly, null *lux* and *cca1* mutant lines produced a smaller number of conidiophores despite increased hyphal biomass amounts.

Inoculation of *Arabidopsis* CRG mutants with the compatible isolate *Hpa*-Maks9 was carried out in the same way as with *Hpa*-Noks1. Reduced sporulation was observed in seven lines: *prr3* (74%), *prr7* (72%), *elf3* (71%), *cca1* (69%), *toc1* (61%), *prr5* (56%) and *phyb* (55%), suggesting the genes are required for compatibility. Enhanced sporulation was observed in three lines (Figure 8C): *kat2* (177%), *det1* (163%) and *lhy* (150%), indicating that these genes play a significant role for basal defence against *Hpa*-Maks9.

*Hpa-*Maks9 hyphal biomass was evaluated for these ten mutant lines: *cca1* (73%), *det1* (70%), *toc1* (66%) and *elf3* (62%) produced less biomass than then control; by contrast, prr7 (155%), *prr5* (141%), *phyb* (138%), *lhy* (137%), *prr3* (133%) and *kat2* (117) had more biomass than that obtained with control Col-0 (Figure 8D). Biomass production in *phyb, prr3, prr7, prr5* were clearly higher than that in control at 3dpi, whereas sporulation was reduced at 7dpi.

Result obtained with *Hpa*-Maks9 on *elf, phyb* and *cca1* mutant lines were similar to those recorded with *Hpa*-Noks1, showing less sporulation than that on controls. Contrastingly, *kat2* and *lhy* mutant lines supported less sporulation, and *toc1* showed more sporulation with *Hpa*-Noks1 infection, giving totally opposite results to those obtained with Maks9. These differences suggest that some mutants might have isolate-specific effects.

As *lhy/toc1* and *cca1/lhy* double mutants, cca1/lhy/toc1 triple mutants were in Ws-0 background, the Ws-compatible isolate *Hpa*-Emco5 were used for both sporulation and biomass production assays. While overexpressors including TOC1ox (113%), LHYox (102%) showed results similar to that obtained from the control, *cca1/lhy* (40%), *lhy/toc1* (34%) and a triple mutant line *cca1/lhy/toc1* (%3) showed statistically significant less sporulation with *Hpa*-Emco5 compared to the Ws control (Figure 8E). *Hpa* biomass was investigated 3 dpi and the results were consistent with the sporulation data (Figure 8F): *lhy/toc1* 46%, *cca1/lhy* 49%, and the triple mutant line *cca1/lhy/toc1* showed the lowest biomass production with 12% (Figure 8F).

## Discussion

Circadian rhythms have long been known to govern various aspects of plant physiology, from growth, differentiation, metabolism and flowering to responses to environmental stresses (Annunziata *et al*., 2018; Luklova *et al*., 2019; Liang *et al*., 2022; Zhu *et al*., 2022). Similarly, microbial microorganisms including fungi and cyanobacteria have their own circadian rhythms that influence processes such as metabolism, nutrient uptake and virulence (Brody, 2019; Valim *et al*., 2022). Previously we showed that light plays a significant role in the development of *Hpa* (Telli *et al*., 2020). In this study, our exploration of the interplay between circadian rhythms and plant-microbe interactions in the *Arabidopsis-Hpa* pathosystem revealed the circadian-related genes (CRGs) *HpaTIM* and *HpaSRR1*. This current study extends our understanding to a biotrophic oomycete pathogen, offering compelling evidence that *Hpa* possesses its own circadian-regulated genes that are necessary for full virulence and provides evidence that circadian rhythms in a biotrophic pathogen can be influenced, directly or indirectly, by CRGs in the plant host.

The discovery of *HpaTIM*, and *HpaSRR1* genes in a plant-obligate oomycete pathogen emphasizes the universal nature of circadian rhythms across diverse organisms. *Drosophila TIM* (Myers *et al*., 1995) plays a central role in entrainment to light-dark (LD) cycles, an adaptation critical for organisms to synchronize with their environment. The rhythmic expression of *HpaTIM* in response to different light conditions indicates its involvement in regulating the pathogen’s circadian clock. Similarly, *SRR1*, or *SENSITIVITY TO RED LIGHT REDUCED 1*, is another key gene found to be essential for circadian regulation (Staiger *et al*., 2003). Its role in influencing various aspects of plant growth and development, as well as its homologues in other organisms, highlights its significance. The presence of *HpaSRR1* and its rhythmic expression further underscores the similarity of biological clocks in both the host and pathogen.

Our recent work demonstrated the efficacy of SS-dsRNA targeting the *Hpa-CesA3* gene, resulting in inhibited spore germination and plant infection (Bilir *et al*., 2019). We used this method to target *HpaTIM* and *HpaSRR1* and the results that both genes *HpaTIM and HpaSRR1* are crucial for pathogen virulence. Interestingly, the mRNA levels of *HpaTIM* and *HpaSRR1* remained unaltered, contrasting with complete suppression of mRNA from positive control *Hpa-Ces3*. Two distinct gene silencing phenomena has been reported: transcriptional and post-transcriptional gene silencing (TGS and PTGS, respectively (Sijen *et al*., 2001). Notably, small RNA studies predominantly implicate PTGS, where they modulate gene expression by base pairing with mRNA targets, leading to degradation or translational inhibition (Saxena *et al*., 2003). Our findings with SS-dsRNAs may be explained by PTGS that inhibits translation of *HpaTIM* and *HpaSRR1* RNA.

One of the most intriguing findings of this study is the correlation of circadian rhythms between *Arabidopsis* and *Hpa*. We observed that the expression patterns of *HpaTIM* and *HpaSRR1* mirrored those of *Arabidopsis* circadian biomarkers *CCA1* and *LHY*. In principle, this correlation could be coincident with no regulatory connection between host and pathogen, due to both organisms’ exposure to the same light regime during infection. On the other hand, we demonstrated that expression of *HpaTIM* and *HpaSRR1* are disrupted by mutations in *CCA1* and *LHY*. Based on this result, we hypothesize that *Hpa* circadian rhythms are coordinated by the *Arabidopsis* clock. It is well documented that host clock can influence rhizosphere microbial community (Hubbard *et al*., 2018; Lu *et al*., 2021; Newman *et al*., 2022). Similarly, the rhizosphere microbial community affects the host clock function (Hubbard *et al*., 2021). These studies suggest there is a bidirectional rhythmic interaction between plants and their rhizomicrobiome (Xu & Dodd, 2022). Investigations with *Drosophila* and its gut microbiome led to the conclusion that microbiome stabilizes circadian rhythm in the host gut to prevent rapid fluctuations with changing environmental conditions (Zhang *et al*., 2023). In addition, the synchronization between the gut microbiome and the host in humans involves intricate crosstalk influenced by diet, lifestyle, and host genetics. This dynamic interaction impacts immune modulation, metabolism and overall health (Thursby & Juge, 2017).

The circadian system can be disturbed transiently or permanently by different factors including light (Telli *et al*., 2020) and temperature (Annunziata *et al*., 2018). Such disturbances are referred to as “circadian dysrhythmia (Bishehsari *et al*., 2020) or “circadian disruption” (Vetter, 2020). Circadian disruption could result in altered microbiome communities and perturbed host metabolism in human health (Bishehsari *et al*., 2020), and changes in global responses in plant immune system (Wang *et al*., 2011). We reported that altering light conditions delayed or inhibited *Hpa* development and sporulation (Telli *et al*., 2020). The data presented in this study represent another step towards defining whether and how plant and pathogen clocks interact, as well as whether circadian dysrhythmia is a factor in plant-pathogen interactions. It will be of interest to assess bidirectional rhythmic interaction between *Hpa* and *Arabidopsis* using gene silencing for *Hpa* and more detailed characterization of *Arabidopsis* CRG mutants.

A recent study on *Arabidopsis* demonstrated that disruption of the plant circadian clock is associated with altered rhythmicity of rhizosphere bacteria and fungi (Newman et al, 2022; (Xu & Dodd, 2022). Here we used *Arabidopsis* single or double clock mutants *cca1 and cca1/lhy* and investigated the expression of *HpaTIM* and *HpaSRR1*. There was an alteration in the circadian expression pattern of these genes on single and double mutants suggesting that the host clock could influence the pathogen clock.

As the influence of the host CRGs on the expression patterns of *HpaTIM* and *HpaSRR1* was clear, we then used 26 singles, 2 double-, 1 triple mutant and also 3 overexpressors (ox) of host clock genes to determine if host clock genes can influence infection and development of the pathogen. We used relevant virulent isolates and measured sporulation and vegetative growth (hyphae). Results of these investigations suggest that CRGs could contribute to compatible interactions as well as basal defence. In addition, double and triple mutants considerably reduced pathogen growth indicating the host CRGs has a major role impact on compatibles interaction. The plant circadian genes impact on large network of defence and development pathways, affecting multiple aspects of biology (Hua, 2013; Luklova *et al*., 2019). It is plausible that access to nutrients by *Hpa* isolates, the amount of host metabolites in the infected tissues and the host redox homeostasis will be altered by some of these CRG mutants resulting in the influence on the pathogen development.

Effector- and PAMP-triggered immunity (ETI and PTI) have been studied in detail in plant-microbe interactions. Control of the *R*-gene mediated defence responses to *Hpa* has been shown to be regulated by CCA1 in *Arabidopsis* (Wang *et al*., 2011). Similarly, PTI against *Pseudomonas syringae* in *Arabidopsis* have been demonstrated to be modulated by the circadian clock (Bhardwaj *et al*., 2011). A recent study on *GIGANTIA* (*GI*) in *Arabidopsis* and the wheat linked the circadian clock to plant susceptibility to pathogens (Kundu & Sahu, 2021), indicating the compatibility may possibly be regulated by the plant’s clock function. Our study on *Arabidopsis* CRG mutants with different virulent *Hpa* isolates clearly indicates the link between host circadian clock and the pathogen in a compatible interaction. Some CRG mutants had different effects on virulence of the two compatible isolates used in this study, suggesting that those isolates might differ in their response to various CRG-regulated pathways that are relevant to *Hpa* virulence. This is another avenue of potential interest in the future.

While we have identified circadian-regulated genes in *Hpa* and possible synchronization with *Arabidopsis* clock genes, the specific mechanisms and the functional implications of this coordination require further investigation. Future research could focus on deciphering the molecular pathways and the specific genes involved in this synchronization, as well as their impact on the development and pathogenicity of the downy mildew pathogen.

## Materials and Methods

### Plant lines, pathogen isolates and their propagation

*H. arabidopsidis* isolates Emoy2, Noks1, Maks9 and Emco5 were maintained on *Arabidopsis* accessions Ws-*eds1 (Parker et al., 1996)* or Col-*rpp4* (Roux *et al*., 2011). Inoculum preparation followed established protocols (Tör *et al*., 2002; Woods-Tör *et al*., 2018)

### Identifying orthologues of two circadian clock genes in *Hpa* genome

Important circadian genes published in model organisms for circadian clock studies were identified through literature. Protein domain-searches through Pfam database (http://pfam.xfam.org) revealed two putative CRGs in *Hpa* genome: *Timeless* (Pfam: PF04821; *Hpa-G810921*, designated *HpaTIM*) and *Sensitive to Red Light Reduced 1 (*Pfam: PF07985; *Hpa-G801448*, designated *HpaSRR1*).

### Time course experiments

Col-*rpp4* seedlings were infected with *Hpa*-Emoy2 as described previously in (Tör *et al*., 2002). Samples were taken every 6h between 4 dpi to 7 dpi from infected seedlings. Total RNA was extracted and analysed with Real-Time PCR. Real-Time PCR analysis used *Hpa-Actin* or *At-Actin* genes as housekeeping genes. The results of samples were analysed by Roche LightCycler 480 Real-Time software program. Each group of experiment had three biological replicas and was repeated thrice. All samples were kept under the 12h Dark/ 12h Light (D/L) cycle, referred to hereafter as “normal”. The inoculated samples were exposed to 4 different light regimes 3 dpi; 1) exposed to constant light between 3 dpi to 4 dpi, then normal 12h L/12h D cycle; 2) exposed to constant dark between 3 dpi to 4 dpi, then normal 12h L/12h D cycle; 3) exposed to constant light 3 dpi until 7 dpi; and 4) exposed to constant dark 3 dpi until 7 dpi. Similarly, in these experiments, samples were taken every 6 hours 4 dpi, and they were analysed with qRT-PCR.

### RNA extraction and gene expression analysis using qRT-PCR

An experiment was designed to confirm the expression pattern of *Arabidopsis* clock genes *CCA1*, *TOC1* and *LHY1*. Uninfected Col-*rpp4* seedlings were used, only sprayed with water, and placed into growth cabinet for a week at 16°C with a photoperiod of 12h dark / 12h light. Samples were taken at 6h intervals 4 dpi and they were analysed using qRT-PCR with the relevant primers (Supplemental Table 1).

To assess dsRNA-mediated gene silencing, gene expression analysis was conducted on four-week-old Ws-*eds1* plants inoculated with a *Hpa*-Emoy2 spore suspension of 5 × 10^4^ spores/mL containing 5 µM siRNA. Six leaves in total were drop-inoculated with a mixture of spores and siRNA (30µl per leaf), with two leaves serving as a biological replicate. The Arabidopsis plants were then placed in a magenta box and subjected to a 12-hour light/12-hour dark regime at 16°C for 3 days.

RNA extraction at different time points after inoculation was carried out. Total RNA extraction protocol by TRIzol (Thermo Fisher) was adapted from (Chomczynski & Sacchi, 2006). Quantitative Real-time PCR (qRT-PCR) was carried out using SensiFAST™ SYBR® No-ROX One-Step Kit (Bioline) and the results were analysed using the 2^−ΔΔCt^ method (Livak & Schmittgen, 2001). Primers for the housekeeping gene for *Arabidopsis* or *Hpa*, *At-Actin or Hpa-Actin*, were included as a control to normalize the results.

A master mix containing; 5 µl of SensiFAST™ SYBR® No-ROX One-Step mix, 2 µl of template (RNA, 20-30ng/µl) from each sample, 0.5 µl of each primer (Supplemental Table 1), 0.01 µl of reverse transcriptase, 0.002 µl RNA-inhibitor was prepared and DEPC water was added to give a final reaction volume of 10 µl. qRT-PCR reactions were carried out in 96-well plates using a Roche Light Cycler Real-Time PCR System. *PCR conditions were as follows,* 45^°^C 10 min, 95^°^C for 2 min, followed by 9 cycles touchdown procedure; 95°C for 5 s; 1^°^C in each annealing step of (68-60) ^°^C for 10 s, 72^°^C 5 s, then 31 cycles of 95^°^C for 5 s, 60^°^C for 10 s, 72^°^C for 5 s.

### Selection of *At*-*CRG*s genes and their mutant lines

*Arabidopsis* TAIR database (https://www.arabidopsis.org) were searched using the “clock gene” keyword and 46 loci matching with 131 distinct gene models were identified. These were then further evaluated for their function and involvement in the circadian rhythm using the available literature. Twenty-six different genes that may play a role in circadian regulation were then selected (Supplemental Table 2).

### Identification of homozygous T-DNA mutants

Totally, 26 singles, 2 double and 1 triple *Arabidopsis* mutant lines along with 3 overexpressors (ox) were selected. Seed of the mutant lines were obtained from NASC (https://arabidopsis.info). To confirm the identity and homozygosity of each mutant, specific primers for each T-DNA line (Supplemental Table 2) were designed using SALK site (http://signal.salk.edu/tdnaprimers.2.html) and were ordered from Sigma (https://www.sigmaaldrich.com/GB/en).Ten seeds from each T-DNA mutant line were sown and seedlings were picked and transferred to pots. DNA was extracted using the REDExtract-N-Amp Tissue PCR Kit protocol (Sigma-R4775) according to the manufacturer’s instruction. DNAs were then used for PCR amplifications. Using this protocol, all lines were screened, and homozygous plants were propagated to obtain seeds for subsequent analyses.

### Sporulation and biomass production assays on T-DNA mutant lines

T-DNA lines were screened with *Hpa*-Maks9, *Hpa*-Noks1 and *Hpa*-Emco5. The samples were taken at 3 dpi and their DNAs extracted for biomass production. DNA was isolated using CTAB method (Doyle, 1987) and the quantitative PCR was performed in a total of 25 µl containing 50ng of gDNA, 12.5 µl of SyberGreen Mastermix (ABI, Carlsbad,California), *Hpa-Actin* or *At-Actin* primers (Supplemental Table 1) and water on a Roche LightCycler 480 device. PCR conditions were as follows, 95^°^C for 4 min, then 10 cycles touchdown of 95^°^C for 30 s, annealing temperature of 65°C, decreasing 1°C every cycle to 56^°^C, and extension at 72°C for 30 s. After 10 cycles of touchdown, a further 25 cycles of 95^°^C for 30 s, 60^°^C for 30 s and 72^°^C for 30 s and a final extension at 72^°^C for 5 min were carried out. Biomass production and sporulation were assessed using quantitative PCR and established protocol as described (Telli *et al*., 2020).

### Application of SS-dsRNAs to pathogen spores and plant inoculations

Silencing of *HpaTIM* and *HpaSRR1* were performed using 30nt-long dsRNA as described previously (Bilir *et al*., 2019). *Hpa-CesA3* were used as a positive control in the silencing experiments. RNA duplexes (Supplemental Table 3) were obtained as synthesised ribonucleotides from Merck.

### Statistical analysis

Statistical analyses were performed using MiniTab Express^TM^ (https://www.minitab.com/en-us/) and GraphPad prism version 10.1.1(GraphPad software, Inc. USA) computer software was used for the statistical analysis. All tests in this study were performed in triplicate. The significant differences among the means were analysed by one-way analysis of variance (ANOVA) complemented by Dunnett’s test at the p < 0.05 level.

### Bioinformatics and phylogenetics

Primer design was performed using Geneious (v10.0) (Kearse *et al*., 2012). We used the EnsemblProtist (Kersey *et al*., 2016) and InterPro (Quevillon *et al*., 2005) databases to identify candidate *Hpa* clock genes. Reciprocal BLASTN and BLASTX (Altschul *et al*., 1997) were used to perform similarity-searches of nucleotide and amino acid sequences, respectively, between *Hpa* clock genes and oomycete and *Arabidopsis* sequences in the UniProt (The UniProt Consortium, 2023) and GenBank (Sayers et a., 2022) databases. To generate multiple sequence alignments against Pfam domains, we used the hmmalign tool from the HMMER package version 3.4 (Eddy, 2011) and profile-HMMs downloaded from Pfam 36.0 (Mistry et al., 2021). Alignments were trimmed, visualised and rendered using JalView (Waterhouse et al. 2009).

Evolutionary histories of protein sequences were inferred by using the maximum-likelihood method and JTT matrix-based model (Jones et al., 1992) conducted in MEGA11 (Tamura et al., 2021). The trees with the highest log likelihoods are shown. Initial tree(s) for the heuristic search were obtained automatically by applying Neighbor-Join and BioNJ algorithms to a matrix of pairwise distances estimated using the JTT model, and then selecting the topology with superior log likelihood value. All positions with less than 95% site coverage were eliminated, i.e., fewer than 5% alignment gaps, missing data, and ambiguous bases were allowed at any position (partial deletion option). Bootstrapping (*i.e.* random sampling with replacement) was performed to generate 1000 bootstrapped trees (Felsenstein, 1985). Phylogenetic trees were rendered and visualised using iTOL (Letunic and Bork, 2021). The trees are drawn to scale, with branch lengths measured in the number of substitutions per site.

## Author contributions

MT and OT planned and designed the research. OT and DG conducted the laboratory work. OT, WJ and DJS performed bioinformatic research, OT, BC-K, YH, JMM, DJS and MT were involved in the analysis of data and wrote the manuscript.

## Data availability statement

The data that support the findings of this study are available from the corresponding author on reasonable request.

---

**Table 1.**
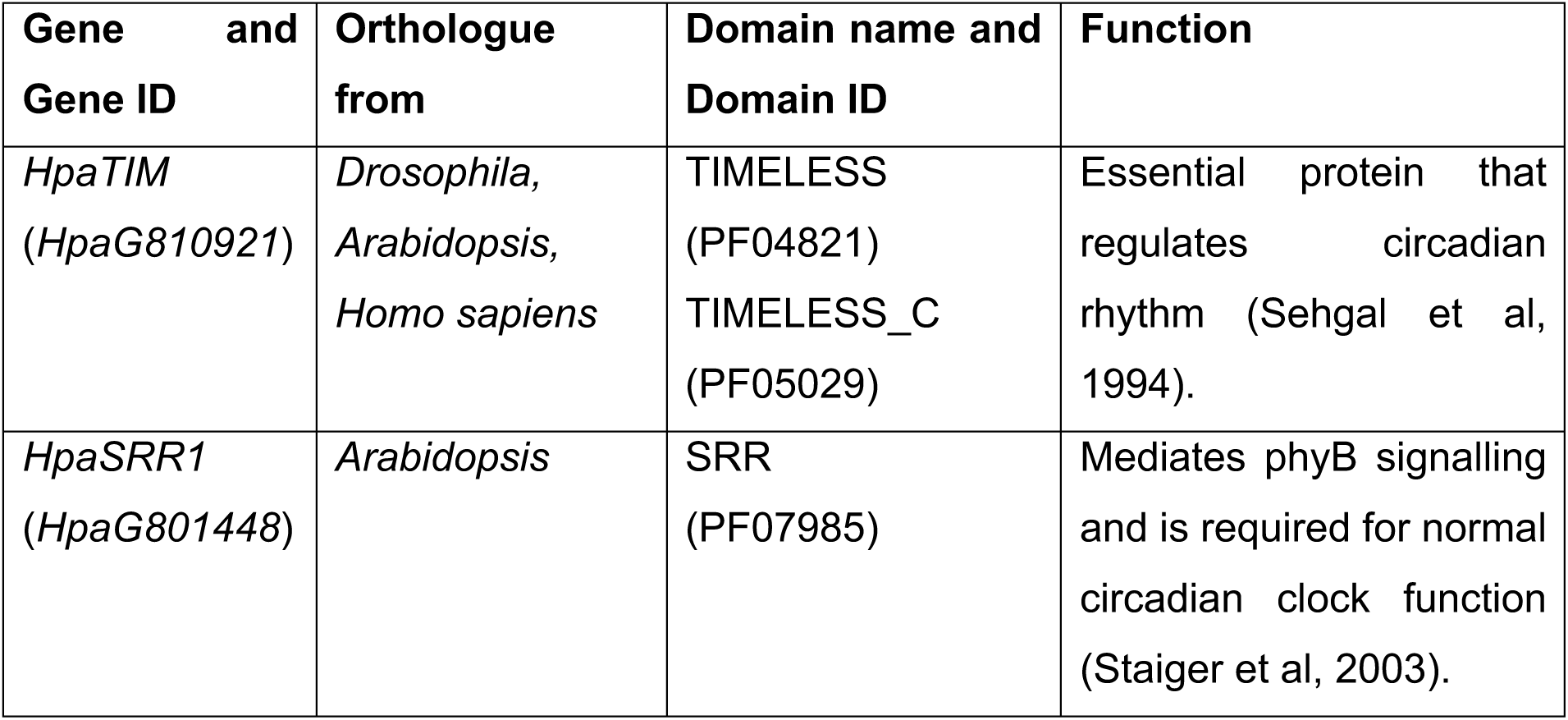
Putative Hpa circadian genes and their orthologues.

| **Gene and Gene ID** | **Orthologue from** | **Domain name and Domain ID** | **Function** |
| --- | --- | --- | --- |
| *HpaTIM*<br>(*HpaG810921*) | *Drosophila*,<br>*Arabidopsis*,<br>*Homo sapiens* | TIMELESS<br>(PF04821)<br>TIMELESS_C<br>(PF05029) | Essential protein that regulates circadian rhythm (Sehgal et al, 1994). |
| *HpaSRR1*<br>(*HpaG801448*) | *Arabidopsis* | SRR<br>(PF07985) | Mediates phyB signalling and is required for normal circadian clock function (Staiger et al, 2003). |

