## Supplementary figures and images for "Synchronization of Circadian Clock Gene Expression in *Arabidopsis* and *Hyaloperonospora arabidopsidis* and its Impact on Host-Pathogen Interactions"

### Supplemental Figure 1

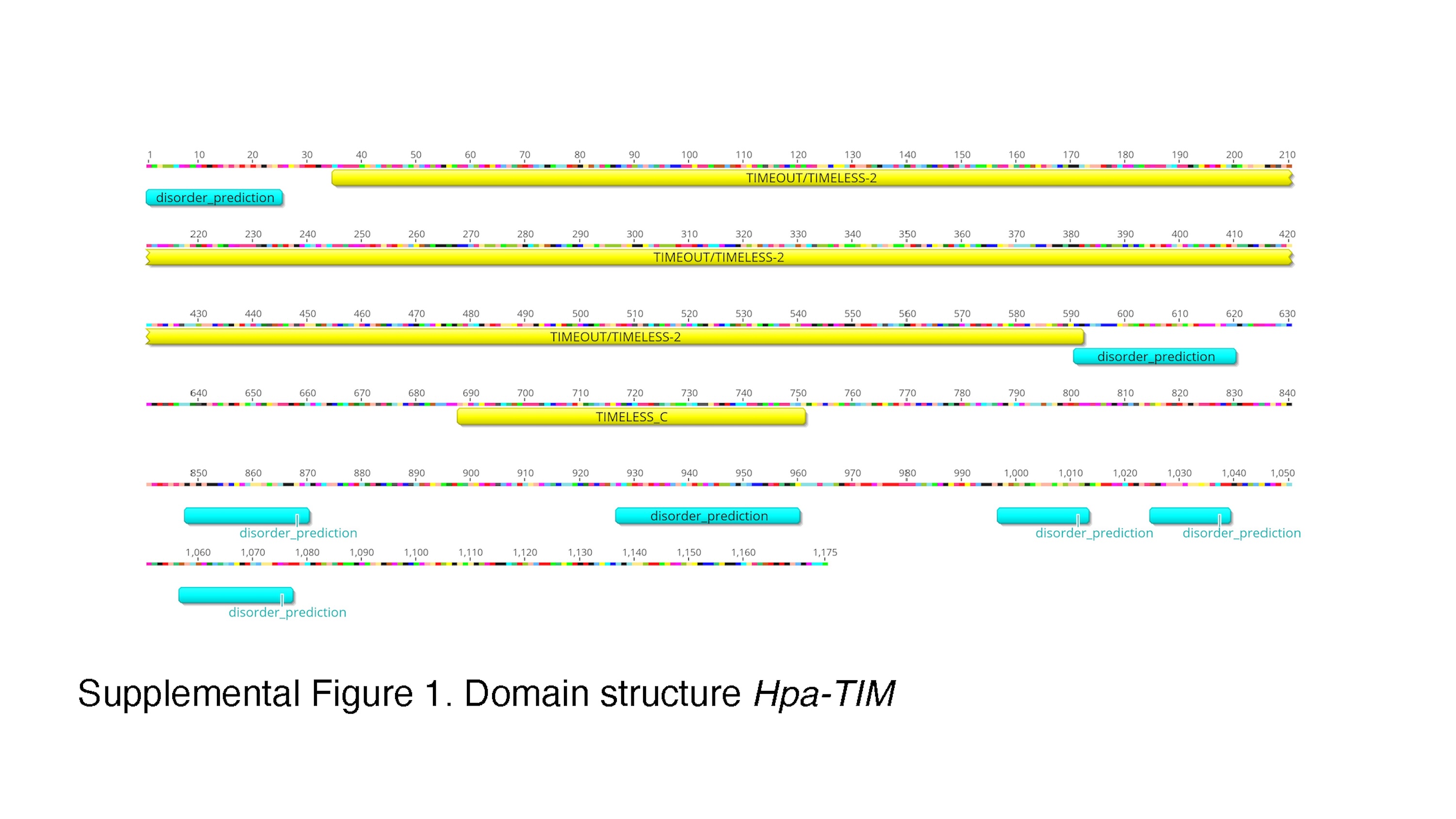

### Supplemental Figure 3

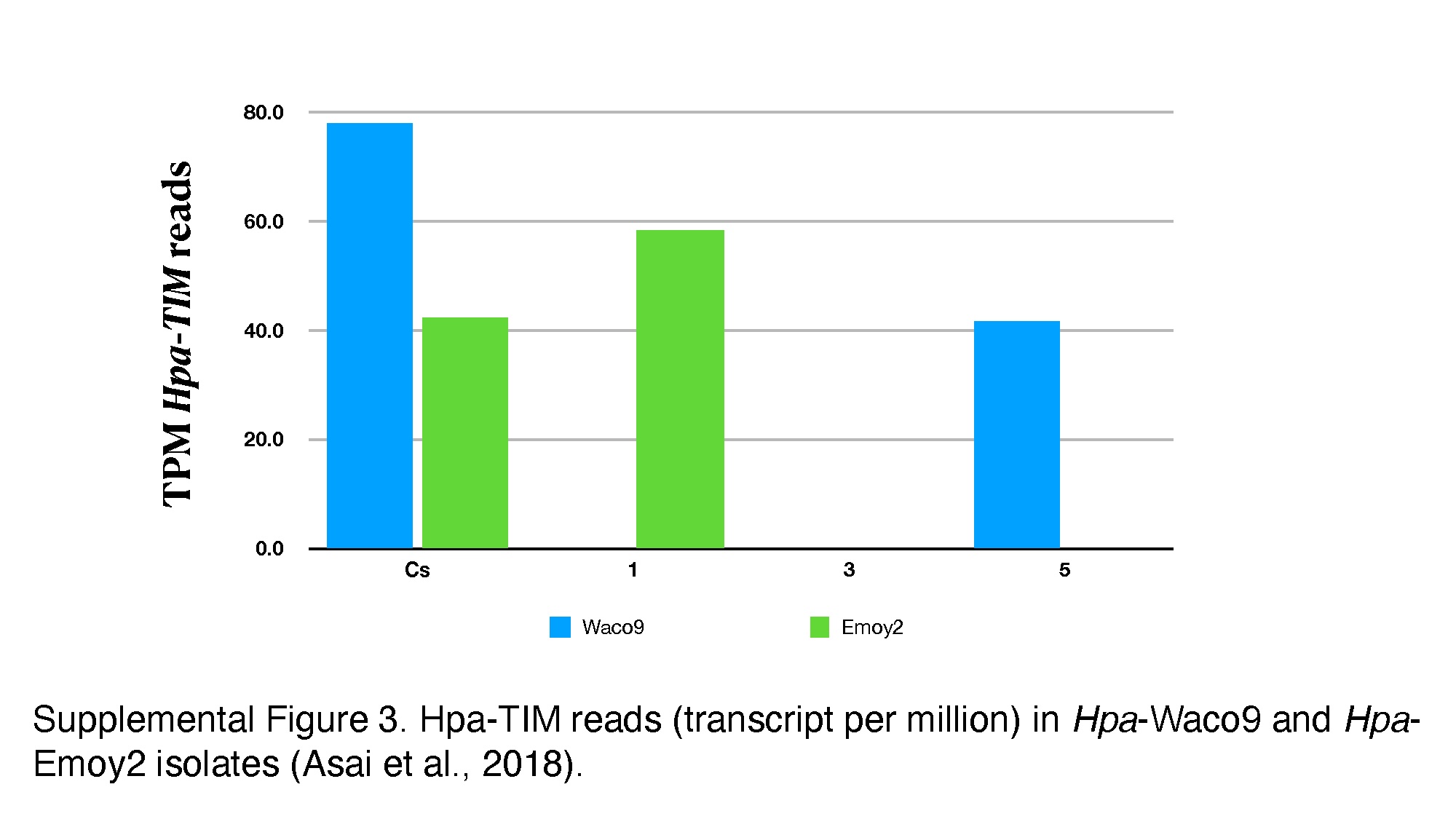

### Supplemental Figure 4

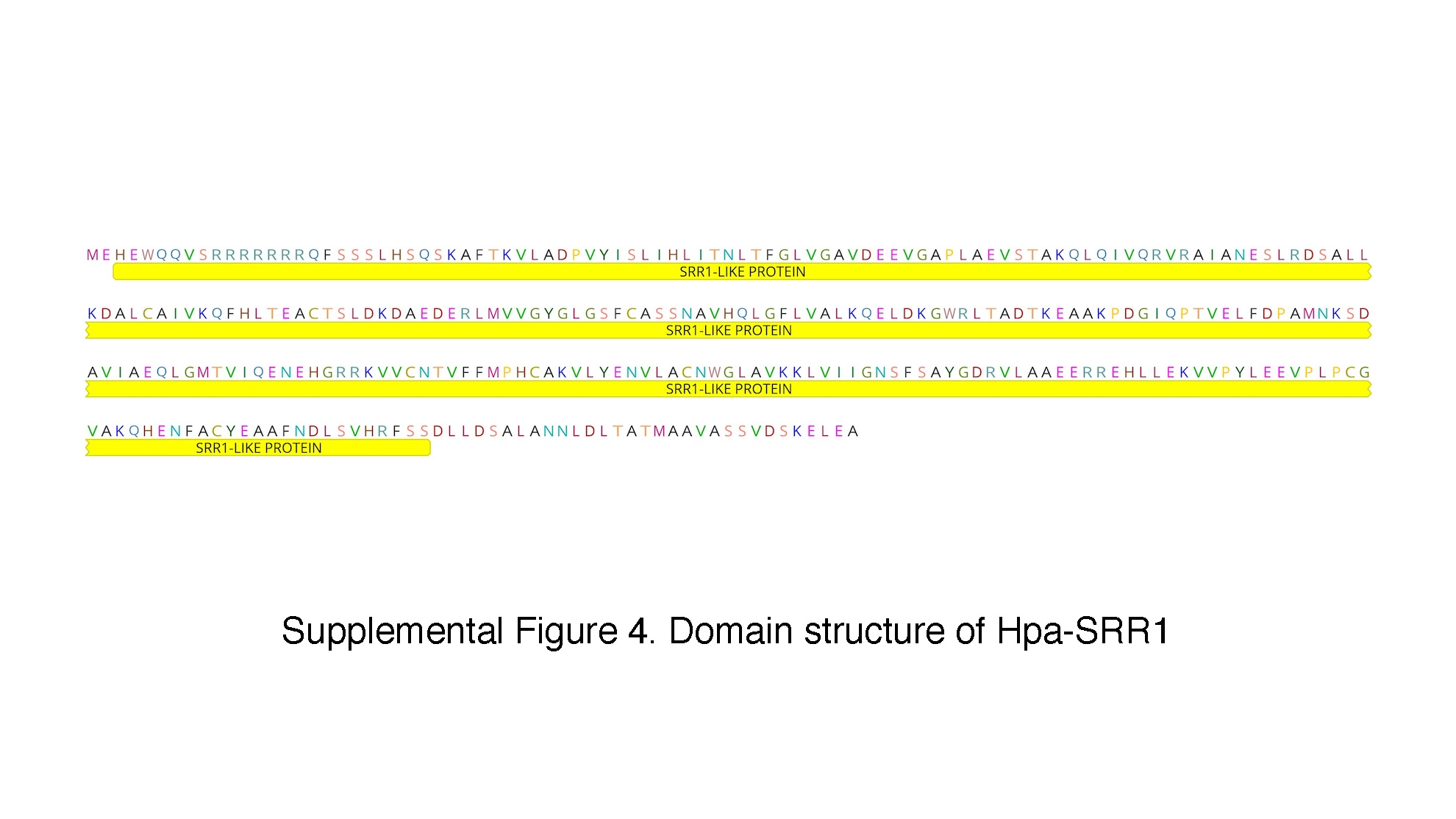

### Supplemental Figure 5

Launch in Jalview

View raw BioJSON Data

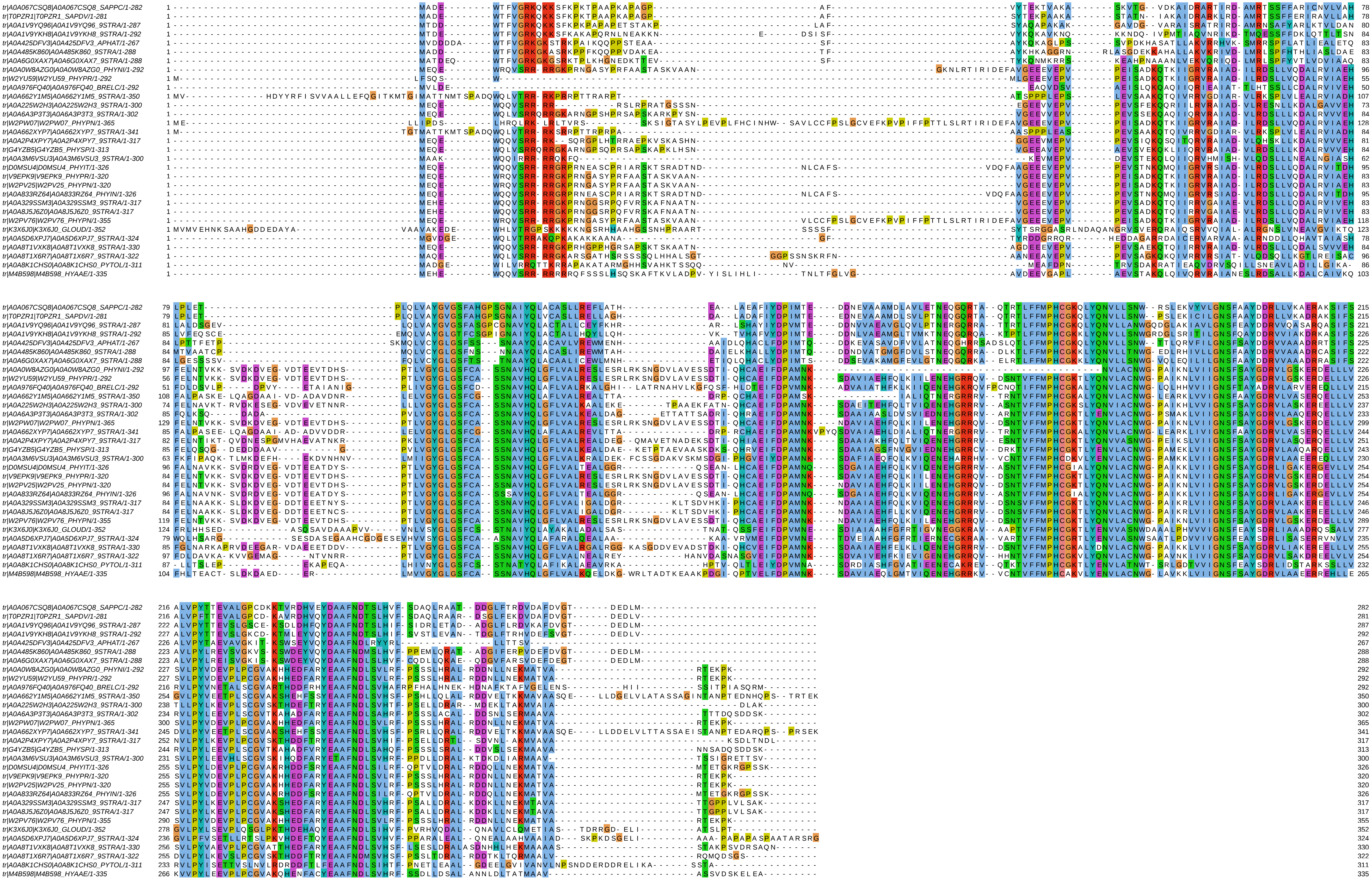
