## Supplemental Figure 2 for "Synchronization of Circadian Clock Gene Expression in *Arabidopsis* and *Hyaloperonospora arabidopsidis* and its Impact on Host-Pathogen Interactions"

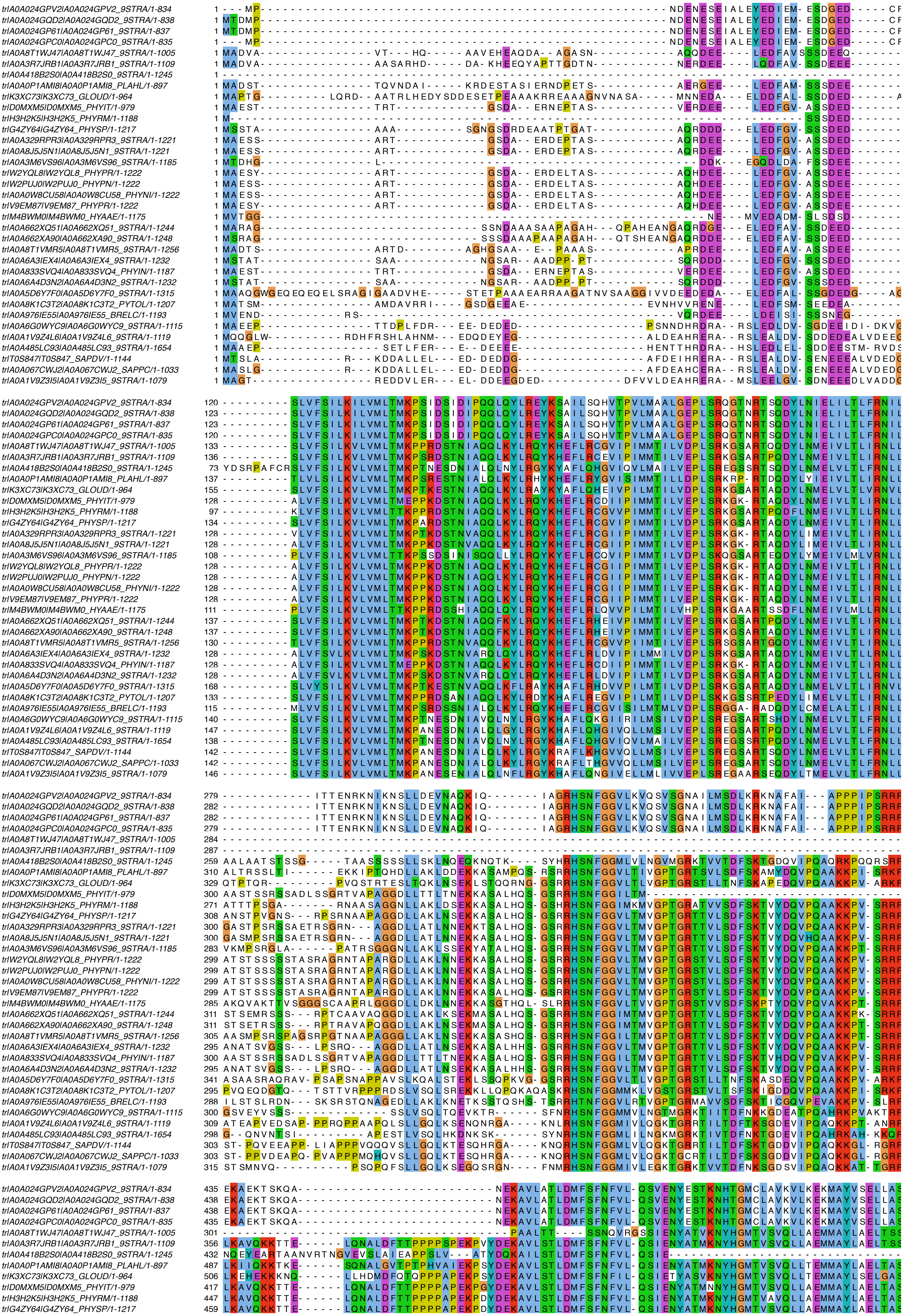

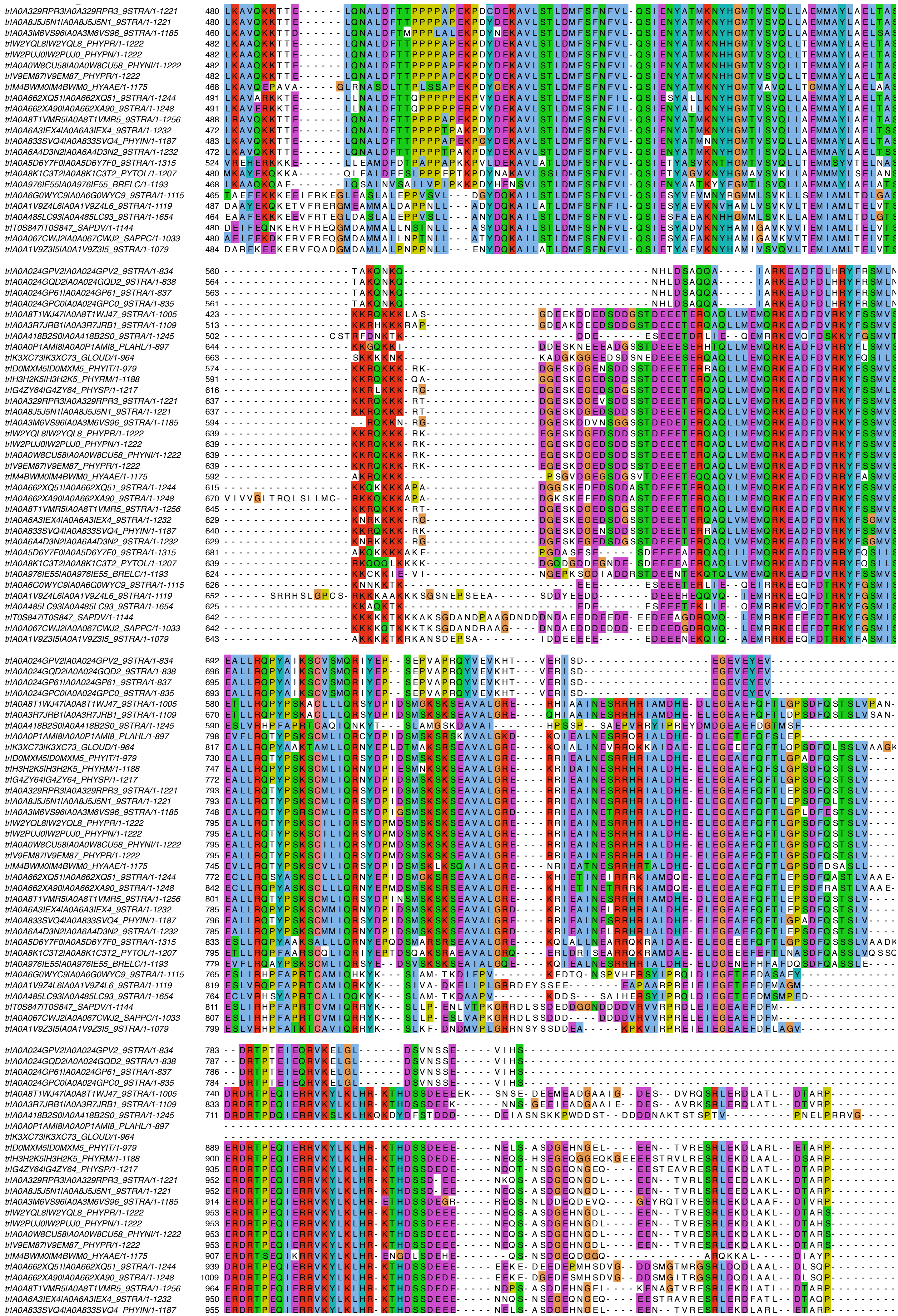

```
tr|A0A6A4D3N2|A0A6A4D3N2_9STRA/1-1232    950 ERDRTPEQIERRVKYLKLHR-KTHDSSDEEE-----NNQS-NSEGEQNGEQ----EESTRAVRESRLERDFATL--DTAQP---------
tr|A0A5D6Y7F0|A0A5D6Y7F0_9STRA/1-1315   1009 DRDRTPEQIERRVKYLKLHR-KTHDSSDEEDERKSADDDY-NDDDERDNGS---ELEDLDAETAARASSD--AR--GSTRP---------
tr|A0A8K1C3T2|A0A8K1C3T2_PYTOL/1-1207    959 DRDRTPEQIERRVKHLKLHR-LTHDSVSEDEDGN--EEDANADDADEDDEDGDNRKPTADPSTEYRLPKGLES---DESDPEE-GLQRAN
tr|A0A976IE55|A0A976IE55_BRELC/1-1193    940 ERDRTPEQIEQRVKFLELQQ-MKHDSLFEDDIG-----------AATDGGQVA-----------DTVEKNLALL--DTARP---------
tr|A0A6G0WYC9|A0A6G0WYC9_9STRA/1-1115    892 DRDRTPEQIERRVKHLKLHK-MDHDSSDEEI-----ES---TGKDPWD-----KDDEESDLVAKDKL---------SIEEPTRVNPDRA-
tr|A0A1V9Z4L6|A0A1V9Z4L6_9STRA/1-1119    945 DRDRTPEQIERRVKHLKLHK-KDHDSSGDED-----------------------------SAADDVI---------AQSSPK--------
tr|A0A485LC93|A0A485LC93_9STRA/1-1654    889 DRDRTPEQIERRVKQLKLHK-KEHDSSSDED-----ESKK-DGKDPWDDS---DDDEEQPSEGTQNM---------PTEAPTRVNPDRG-
tr|T0S847|T0S847_SAPDV/1-1144            945 DRDRTPEQIERRVKHLKLHK-KSHDDDDDDD-----------------------NASDAMSATDDNV---------APRSPKP-------
tr|A0A067CWJ2|A0A067CWJ2_SAPPC/1-1033    937 DRDRTPDQIERRVKHLKLHK-KTHDDDDDDD-----DDGG-------------DDDDDAMSATDDNV---------APRSPKP-------
tr|A0A1V9Z3I5|A0A1V9Z3I5_9STRA/1-1079    927 DRDRTPEQIERRVRHLKLHK-KEHDLSDESD-----EE----------------------ATQNDTP---------AEVIPTT-------

tr|A0A024GPV2|A0A024GPV2_9STRA/1-834         ------------------------------------------------------------------------------------------
tr|A0A024GQD2|A0A024GQD2_9STRA/1-838         ------------------------------------------------------------------------------------------
tr|A0A024GP61|A0A024GP61_9STRA/1-837         ------------------------------------------------------------------------------------------
tr|A0A024GPC0|A0A024GPC0_9STRA/1-835         ------------------------------------------------------------------------------------------
tr|A0A8T1WJ47|A0A8T1WJ47_9STRA/1-1005    861 ESDGARLSLNEKIY------------DSTAGAQ--------------GDGT---------------------------------KESQTQ
tr|A0A3R7JRB1|A0A3R7JRB1_9STRA/1-1109    952 KSDIARHAPSEKRY------------DSTVSVQ--------------GGDA---------------------------------NESQTQ
tr|A0A418B2S0|A0A418B2S0_9STRA/1-1245    846 AKN--DDSDDDDLDMLL------SRGSSSTWSPEQT---------------------------KRRR-----------------------
tr|A0A0P1AMI8|A0A0P1AMI8_PLAHL/1-897     880 -------------------------------------------------------------------------------------SNTNQ
tr|K3XC73|K3XC73_GLOUD/1-964                 ------------------------------------------------------------------------------------------
tr|D0MXM5|D0MXM5_PHYIT/1-979                 ------------------------------------------------------------------------------------------
tr|H3H2K5|H3H2K5_PHYRM/1-1188           1019 ASN---AADDDEMS------------GKSPTEDVSFTIP------PVSDA----------------------------------NAIQSQ
tr|G4ZY64|G4ZY64_PHYSP/1-1217           1051 ASN---AVHDEEMN----------AEDSSTEVQPSK---------TLHD-----------------------------------DASQTQ
tr|A0A329RPR3|A0A329RPR3_9STRA/1-1221   1065 VLD---ATPDKEMN------------DTSPTVEASK-----------ND--------------------------------------VEQ
tr|A0A8J5J5N1|A0A8J5J5N1_9STRA/1-1221   1065 VLD---AAPDKEMN------------DTSPTVEASK-----------ND--------------------------------------VEQ
tr|A0A3M6VS96|A0A3M6VS96_9STRA/1-1185   1028 VSD---AVSDEEMN------------DAHPAGDLSNEVSS----TNVNDAL---------------------------------DKSYSQ
tr|W2YQL8|W2YQL8_PHYPR/1-1222           1064 ASD---AIPDAEMN------------GTSPTVNSSK-----------NDDT---------------------------------DESQTQ
tr|W2PUJ0|W2PUJ0_PHYPN/1-1222           1064 ASD---AIPDAEMN------------GTSPTVNSSK-----------NDDT---------------------------------DESQTQ
tr|A0A0W8CU58|A0A0W8CU58_PHYNI/1-1222   1064 ASD---AIPDAEMN------------GTSPTVNSSK-----------NDDT---------------------------------DESQTQ
tr|V9EM87|V9EM87_PHYPR/1-1222           1064 ASD---AIPDAEMN------------GTSPTVNSSK-----------NDDT---------------------------------DESQTQ
tr|M4BWM0|M4BWM0_HYAAE/1-1175           1007 VTGSSGAVPDDEMN------------DASLHAESSSETT-----PTLNDDA---------------------------------HKGQCK
tr|A0A662XQ51|A0A662XQ51_9STRA/1-1244   1063 VTEDSEAVRDQQMN----------------------------------------------------------------------GEDASD
tr|A0A662XA90|A0A662XA90_9STRA/1-1248   1133 VTEDLEAVRDQHMS----------------------------------------------------------------------GEDVSD
tr|A0A8T1VMR5|A0A8T1VMR5_9STRA/1-1256   1080 ASD---AVPDDEMN------------VSTPAGDTSNEIS-----TKSNDNA---------------------------------GENESQ
tr|A0A6A3IEX4|A0A6A3IEX4_9STRA/1-1232   1065 ------TVPNEEVN-----------DDSATEVQTS----------TAND-----------------------------------NESQSQ
tr|A0A833SVQ4|A0A833SVQ4_PHYIN/1-1187   1064 -GD---AVPDEEIN------------GTSPTADSSK-----------NEDA---------------------------------NESQSQ
tr|A0A6A4D3N2|A0A6A4D3N2_9STRA/1-1232   1065 ------TVPNEEVN-----------DDSATEVQTS----------TAND-----------------------------------NESQSQ
tr|A0A5D6Y7F0|A0A5D6Y7F0_9STRA/1-1315   1125 AVD-RDGATDEPMDGVP------GDSDSLAAGALPG-VS-----GAESDEV---------------------------------GDGDAQ
tr|A0A8K1C3T2|A0A8K1C3T2_PYTOL/1-1207   1091 --------EMEEMNAGS------VENSSSTGAPDEGGIEET--------------------------Q-----P----------FESEEQ
tr|A0A976IE55|A0A976IE55_BRELC/1-1193   1042 VINGA----FREMTCMS--------------------VK-----VNLRDVD---------------------------------KESQPY
tr|A0A6G0WYC9|A0A6G0WYC9_9STRA/1-1115   1032 TSR-DDNSDDDELDSLSIA-------ASTQAQPDEA-------------ASADENK-------KRPR-----------------------
tr|A0A1V9Z4L6|A0A1V9Z4L6_9STRA/1-1119   1032 RSL--PDDAQDEGGA-----------DSSEDAPVAAAIV---------------------------------------------------
tr|A0A485LC93|A0A485LC93_9STRA/1-1654   1039 RSQEDDDDSEDEFDTLAST--MTTQQESFEETMDDS-IEDTIEDDSVEDSVADESKAEEGSSPKRPRLGSAEAPNEESENY---TFEQSY
tr|T0S847|T0S847_SAPDV/1-1144           1039 RSHDDDDDDDDEGGDDSLLA------PTSENAPSSP------------NATAD---------RKRSR-----------------------
tr|A0A067CWJ2|A0A067CWJ2_SAPPC/1-1033        ------------------------------------------------------------------------------------------
tr|A0A1V9Z3I5|A0A1V9Z3I5_9STRA/1-1079   1021 -----QDDSIEESAIVS---------DDTERVPT--------------------------------------------------------

tr|A0A024GPV2|A0A024GPV2_9STRA/1-834         ------------------------------------------------------------------------------------------
tr|A0A024GQD2|A0A024GQD2_9STRA/1-838         ------------------------------------------------------------------------------------------
tr|A0A024GP61|A0A024GP61_9STRA/1-837         ------------------------------------------------------------------------------------------
tr|A0A024GPC0|A0A024GPC0_9STRA/1-835         ------------------------------------------------------------------------------------------
tr|A0A8T1WJ47|A0A8T1WJ47_9STRA/1-1005    900 ------------------------------EQS------------------------------SGQ-TEF--------------------
tr|A0A3R7JRB1|A0A3R7JRB1_9STRA/1-1109    991 ------------------------------EDS------------------------------SGQ-TELAA------------------
tr|A0A418B2S0|A0A418B2S0_9STRA/1-1245    889 ------------------ASKRTR------HDD------------------------------ESRVADEAQQSE---------------
tr|A0A0P1AMI8|A0A0P1AMI8_PLAHL/1-897     887 ------------------------------------------------------------------------------------------
tr|K3XC73|K3XC73_GLOUD/1-964                 ------------------------------------------------------------------------------------------
tr|D0MXM5|D0MXM5_PHYIT/1-979                 ------------------------------------------------------------------------------------------
tr|H3H2K5|H3H2K5_PHYRM/1-1188           1062 ------------------------------EDSLD-------------------------ARGEGQ-SEQSTDAE---------------
tr|G4ZY64|G4ZY64_PHYSP/1-1217           1092 ------------------------------EDGIR--------------------------------EEQSTEAE---------------
tr|A0A329RPR3|A0A329RPR3_9STRA/1-1221   1099 ------------------------------EDSLD-------------------------SRSQDP-TEQSS------------------
tr|A0A8J5J5N1|A0A8J5J5N1_9STRA/1-1221   1099 ------------------------------ENSLD-------------------------SRSQDP-TEQSS------------------
tr|A0A3M6VS96|A0A3M6VS96_9STRA/1-1185   1074 ------------------------------QDIQH-------------------------SCLEHH-IEQPTKA----------------
tr|W2YQL8|W2YQL8_PHYPR/1-1222           1103 ------------------------------EDNLD-------------------------SHPL---TEQSSEAQ---------------
tr|W2PUJ0|W2PUJ0_PHYPN/1-1222           1103 ------------------------------EDNLD-------------------------SHPL---TEQSSEAQ---------------
tr|A0A0W8CU58|A0A0W8CU58_PHYNI/1-1222   1103 ------------------------------EDNLD-------------------------SHPL---TEQSSEAQ---------------
tr|V9EM87|V9EM87_PHYPR/1-1222           1103 ------------------------------EDNLD-------------------------SHPL---TEQSSEAQ---------------
tr|M4BWM0|M4BWM0_HYAAE/1-1175           1053 ------------------------------ADSIH-------------------------SRHEYQ-IRQPADRQ---------------
tr|A0A662XQ51|A0A662XQ51_9STRA/1-1244   1091 ------------------------------EDEVT------------------------QLREAQ-AEPST-AQ----------------
tr|A0A662XA90|A0A662XA90_9STRA/1-1248   1161 ------------------------------EDEAM------------------------QLREAQ-VEPST--------------------
tr|A0A8T1VMR5|A0A8T1VMR5_9STRA/1-1256   1130 ------------------------------EDSLN-------------------------SRPEDQ-TEQSTEAQ---------------
tr|A0A6A3IEX4|A0A6A3IEX4_9STRA/1-1232   1101 ------------------------------EDSLD-------------------------IRRESQ-TEQPTEAE---------------
tr|A0A833SVQ4|A0A833SVQ4_PHYIN/1-1187   1100 ------------------------------QDQ----------------------------------TEQSSDAL---------------
tr|A0A6A4D3N2|A0A6A4D3N2_9STRA/1-1232   1101 ------------------------------EDSLD-------------------------IRRESQ-TEQPTEAE---------------
tr|A0A5D6Y7F0|A0A5D6Y7F0_9STRA/1-1315   1179 ------------------------------EDDAG-------------------------VTHVSDRDEPTFEGE---------------
tr|A0A8K1C3T2|A0A8K1C3T2_PYTOL/1-1207   1134 ------------------------------ESEEQ------------------------TASSATQ------------------------
tr|A0A976IE55|A0A976IE55_BRELC/1-1193   1078 ------------------------------ERE-----------------------------SQDQ-LEFPVKAH----------------
tr|A0A6G0WYC9|A0A6G0WYC9_9STRA/1-1115   1071 ------------------------------------------------------------------------------------------
tr|A0A1V9Z4L6|A0A1V9Z4L6_9STRA/1-1119        ------------------------------------------------------------------------------------------
tr|A0A485LC93|A0A485LC93_9STRA/1-1654   1213 ---------PPKGRTKPKASKAGR-----SDNQHGN-------DGDAAESSKPTSPALKHEVAKSPVKEKPKKKK---------------
tr|T0S847|T0S847_SAPDV/1-1144           1081 -----------------------------SNDE---------------------------------------------------------
tr|A0A067CWJ2|A0A067CWJ2_SAPPC/1-1033   1026 ------------------------------------------------------------------------------------------
tr|A0A1V9Z3I5|A0A1V9Z3I5_9STRA/1-1079        ------------------------------------------------------------------------------------------

tr|A0A024GPV2|A0A024GPV2_9STRA/1-834         ------------------------------------------------------------------------------------------
tr|A0A024GQD2|A0A024GQD2_9STRA/1-838         ------------------------------------------------------------------------------------------
tr|A0A024GP61|A0A024GP61_9STRA/1-837         ------------------------------------------------------------------------------------------
tr|A0A024GPC0|A0A024GPC0_9STRA/1-835         ------------------------------------------------------------------------------------------
tr|A0A8T1WJ47|A0A8T1WJ47_9STRA/1-1005    934 ----------EEPT-------------TADIVI---------------AEMEATT-----LNDTH------AESVVAGA-ASGVQSLKRA
tr|A0A3R7JRB1|A0A3R7JRB1_9STRA/1-1109   1034 ----------EEPT-------------TTDVVT---------------ADIEATT-----LNSTQ------VEPV--DD-ASGVQSLKRV
tr|A0A418B2S0|A0A418B2S0_9STRA/1-1245    981 AKQAKEELDAKHADGA----EPAHA--GPEQLEKFQKQTKERLLKAKEAKRQAEL-----LRQAQERQQEEMKAKHLADEAASADDAKRF
tr|A0A0P1AMI8|A0A0P1AMI8_PLAHL/1-897         ------------------------------------------------------------------------------------------
tr|K3XC73|K3XC73_GLOUD/1-964                 ------------------------------------------------------------------------------------------
tr|D0MXM5|D0MXM5_PHYIT/1-979                 ------------------------------------------------------------------------------------------
tr|H3H2K5|H3H2K5_PHYRM/1-1188           1112 ADGAAMETDAEH-----------------ESPS---------------VAMEAAS-----LNDDQ-------EQ------NDAAQSLKRA
tr|G4ZY64|G4ZY64_PHYSP/1-1217           1131 AAEDSAMTGVELPD-------------ATEAVT---------------AEMEVAN-----LNGSQ-------EPTNPESDTADVQSLKRG
tr|A0A329RPR3|A0A329RPR3_9STRA/1-1221   1142 VDGANVGSGNEAED-------------T-EMVA---------------AEMEAAK-----INGSQ-------DDVALE---ADAQSLKRG
tr|A0A8J5J5N1|A0A8J5J5N1_9STRA/1-1221   1142 VDGANVGSGNEAED-------------T-EMVA---------------AEMEAAK-----INGSQ-------DDVALE---ADAQSLKRG
tr|A0A3M6VS96|A0A3M6VS96_9STRA/1-1185   1110 PDSADAGSG----D-------------GPETVI---------------AEMEVAT-----ITDSR-------DDVAADD-AADAQPHKRV
tr|W2YQL8|W2YQL8_PHYPR/1-1222           1140 INGANSESGNEAED-------------T-EMVT---------------ADMDAAK-----INGSQ-------VDVVLEE-AADAQSLKRS
tr|W2PUJ0|W2PUJ0_PHYPN/1-1222           1140 INGANSESGNEAED-------------T-EMVT---------------ADMDAAK-----INGSQ-------VDVVLEE-AADAQSLKRS
tr|A0A0W8CU58|A0A0W8CU58_PHYNI/1-1222   1140 INGANSESGNEAED-------------T-EMVT---------------ADMDAAK-----INGSQ-------VDVVLEE-AADAQSLKRS
tr|V9EM87|V9EM87_PHYPR/1-1222           1140 INGANSESGNEAED-------------T-EMVT---------------ADMDAAK-----INGSQ-------VDVVLEE-AADAQSLKRS
tr|M4BWM0|M4BWM0_HYAAE/1-1175           1091 VSTAHEGAKVESTD-------------GPATVT---------------AGMDAAN-----IDGSQ-------DDVIPVD--ADAQPNKRA
tr|A0A662XQ51|A0A662XQ51_9STRA/1-1244   1155 GAEAEEQTGVEERT-------------PTDAVT---------------TEMEAAT-----LSGSQ------QESTVMVAEVADANPLKRS
tr|A0A662XA90|A0A662XA90_9STRA/1-1248   1177 -----------------------------DAVA---------------TEMEAAT-----LNRSQ------HEPAVLVAEVADTNPLKRS
tr|A0A8T1VMR5|A0A8T1VMR5_9STRA/1-1256   1179 TTGSNAETSTEPAD-------------T-EMVT---------------ADMEAAK-----INGSQ-------EDAVLEE-AADAQSLKRG
tr|A0A6A3IEX4|A0A6A3IEX4_9STRA/1-1232   1148 AAEDSAMTGDELPN-------------A-ETVA---------------AETEAAA-----LNVSQ-------ELENPDGGAADVQSLKRS
tr|A0A833SVQ4|A0A833SVQ4_PHYIN/1-1187   1128 ---ANVVAG----D-------------T-EMVT---------------AEMEAAK-----IAGSE-------ENVMLGE-AADVQPLKRG
tr|A0A6A4D3N2|A0A6A4D3N2_9STRA/1-1232   1148 AAEDSAMTGDELPN-------------A-ETVA---------------AETEAAA-----LNVSQ-------ELENPDGGAADVQSLKRS
tr|A0A5D6Y7F0|A0A5D6Y7F0_9STRA/1-1315   1229 LSAANGVDSTEAVD---------------DAVMDG------------ETELEAAA-----V-----------ATAAAGDEEAATPSNKRG
tr|A0A8K1C3T2|A0A8K1C3T2_PYTOL/1-1207   1146 -----------------------------------------------ESDGESER--------------------------------KRF
tr|A0A976IE55|A0A976IE55_BRELC/1-1193   1122 -----------------------------DTFTKKT---------IIETSYEAGTSKINLINVPRT---HGSPDVDVVEEATNTESHKRG
tr|A0A6G0WYC9|A0A6G0WYC9_9STRA/1-1115   1075 ---------------------------------------------------------------------------------DTAESPKRL
tr|A0A1V9Z4L6|A0A1V9Z4L6_9STRA/1-1119   1058 --------------------APLHA------------------------------------------------------------DPKRF
tr|A0A485LC93|A0A485LC93_9STRA/1-1654   1350 LREKQKHDDASSTDEKPEKDVPVKS--NPGQIEKFQQQTKERLLRAKEQKRQAEL-----LKQEQLKRDEDEKLKHKEAEADAAEESKRF
tr|T0S847|T0S847_SAPDV/1-1144           1101 -------------------------------------------------------------------------------------DAKRF
tr|A0A067CWJ2|A0A067CWJ2_SAPPC/1-1033        ------------------------------------------------------------------------------------------
tr|A0A1V9Z3I5|A0A1V9Z3I5_9STRA/1-1079   1041 ---------------------------------------------------------------------------------------KRF

tr|A0A024GPV2|A0A024GPV2_9STRA/1-834         ------------------------------------------------------------------------------------------
tr|A0A024GQD2|A0A024GQD2_9STRA/1-838         ------------------------------------------------------------------------------------------
```

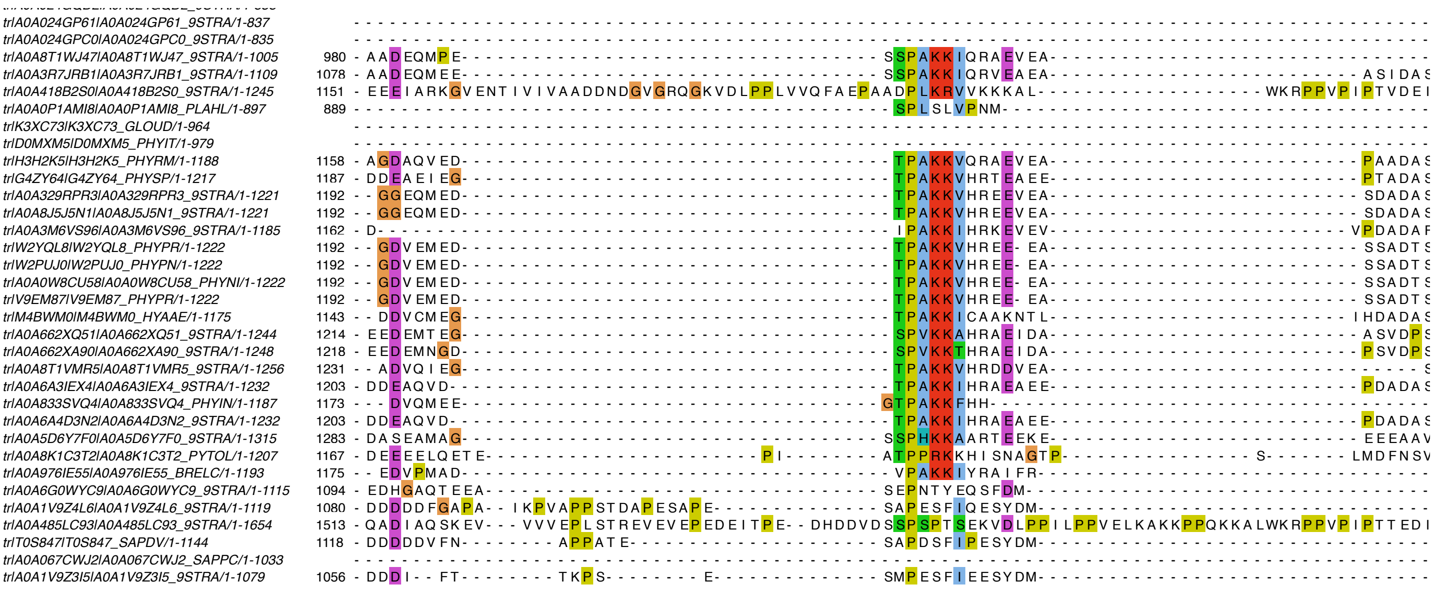

tr|A0A024GP61|A0A024GP61_9STRA/1-837
tr|A0A024GPC0|A0A024GPC0_9STRA/1-835
tr|A0A8T1WJ47|A0A8T1WJ47_9STRA/1-1005 980
tr|A0A3R7JRB1|A0A3R7JRB1_9STRA/1-1109 1078
tr|A0A418B2S0|A0A418B2S0_9STRA/1-1245 1151
tr|A0A0P1AMI8|A0A0P1AMI8_PLAHL/1-897 889
tr|K3XC73|K3XC73_GLOUD/1-964
tr|D0MXM5|D0MXM5_PHYIT/1-979
tr|H3H2K5|H3H2K5_PHYRM/1-1188 1158
tr|G4ZY64|G4ZY64_PHYSP/1-1217 1187
tr|A0A329RPR3|A0A329RPR3_9STRA/1-1221 1192
tr|A0A8J5J5N1|A0A8J5J5N1_9STRA/1-1221 1192
tr|A0A3M6VS96|A0A3M6VS96_9STRA/1-1185 1162
tr|W2YQL8|W2YQL8_PHYPR/1-1222 1192
tr|W2PUJ0|W2PUJ0_PHYPN/1-1222 1192
tr|A0A0W8CU58|A0A0W8CU58_PHYNI/1-1222 1192
tr|V9EM87|V9EM87_PHYPR/1-1222 1192
tr|M4BWM0|M4BWM0_HYAAE/1-1175 1143
tr|A0A662XQ51|A0A662XQ51_9STRA/1-1244 1214
tr|A0A662XA90|A0A662XA90_9STRA/1-1248 1218
tr|A0A8T1VMR5|A0A8T1VMR5_9STRA/1-1256 1231
tr|A0A6A3IEX4|A0A6A3IEX4_9STRA/1-1232 1203
tr|A0A833SVQ4|A0A833SVQ4_PHYIN/1-1187 1173
tr|A0A6A4D3N2|A0A6A4D3N2_9STRA/1-1232 1203
tr|A0A5D6Y7F0|A0A5D6Y7F0_9STRA/1-1315 1283
tr|A0A8K1C3T2|A0A8K1C3T2_PYTOL/1-1207 1167
tr|A0A976IE55|A0A976IE55_BRELC/1-1193 1175
tr|A0A6G0WYC9|A0A6G0WYC9_9STRA/1-1115 1094
tr|A0A1V9Z4L6|A0A1V9Z4L6_9STRA/1-1119 1080
tr|A0A485LC93|A0A485LC93_9STRA/1-1654 1513
tr|T0S847|T0S847_SAPDV/1-1144 1118
tr|A0A067CWJ2|A0A067CWJ2_SAPPC/1-1033
tr|A0A1V9Z3I5|A0A1V9Z3I5_9STRA/1-1079 1056
