## Supplemental Table 1 for "Synchronization of Circadian Clock Gene Expression in *Arabidopsis* and *Hyaloperonospora arabidopsidis* and its Impact on Host-Pathogen Interactions"

| **Supplemental Table 1.**  Primers used in the analysis of *Hpa* and *Arabidopsis* clock genes and *Hpa* gene silencing experiments. | | |
| --- | --- | --- |
| **Gene** | **Forward Primer (5’-3’)** | **Reverse Primer (5’-3’)** |
| *HPATIM* | CTTCAATCTTCTGCCATGCTC | TACCCTTCATGCATGGTTAGC |
| *HPASRR1* | CGTTGGCTATGGACTCGGAA | TCTTTAGTGTCGGCGGTCAG |
| *HPA-Actin* | TGGCTGAAGGGTACGTGATT | TCCGGCTTAGCATCGTACTT |
| *At-CCA1* | TACCCTTCATGCATGGTTAGC | CTTCAATCTTCTGCCATGCTC |
| *At-LHY1* | TGTTTGTGTATGCTACTTGTGGT | CAACGAGCGCCTTCAAGTTT |
| *At-TOC1* | AGAGGGACCGAGCAATACCA | CCATTGATGTGGGAAGCGTG |
| *At-Actin* | AGCATCTGGTCTGCGAGTTC | ACGGATTTAATGACACAATGGC |
| *Hpa-CesA3* | TGTCCGGAACTACTACGAGC | CCGACATCTTGTACTTGCGG |
